# Decryption of sequence, structure, and functional features of SINE repeat elements in SINEUP non-coding RNA–mediated post-transcriptional gene regulation

**DOI:** 10.1101/2023.05.22.541671

**Authors:** Harshita Sharma, Matthew N Z Valentine, Naoko Toki, Hiromi Sueki Nishiyori, Stefano Gustincich, Hazuki Takahashi, Piero Carninci

**Affiliations:** Laboratory for Transcriptome Technology, RIKEN Center for Integrative Medical Sciences, Yokohama, Kanagawa, 230-0045, Japan; Department of Neuroscience and Brain Technologies, Istituto Italiano di Tecnologia, Genova, Italy; Human Technopole, Milan, 20157, Italy

## Abstract

RNA structure folding largely influences RNA regulation by providing flexibility and functional diversity. In silico and in vitro analyses are limited in their ability to capture the intricate relationships between dynamic RNA structure and RNA functional diversity present in the cell. Here, we investigate sequence, structure and functional features of mouse and human SINE-transcribed retrotransposons embedded in SINEUPs long non-coding RNAs, which positively regulate target gene expression post-transcriptionally. In-cell secondary structure probing reveals that functional SINEs-derived RNAs contain conserved short structure motifs essential for SINEUP-induced translation enhancement. We show that SINE RNA structure dynamically changes between the nucleus and cytoplasm and is associated with compartment-specific binding to RBP and related functions. Moreover, RNA–RNA interaction analysis shows that the SINE-derived RNAs interact directly with ribosomal RNAs, suggesting a mechanism of translation regulation. We further predict the architecture of 18 SINE RNAs in three dimensions guided by experimental secondary structure data. Overall, we demonstrate that the conservation of short key features involved in interactions with RBPs and ribosomal RNA drives the convergent function of evolutionarily distant SINE-transcribed RNAs.

## Introduction

In long non-coding RNAs (lncRNAs), evolutionary selection takes place not only at the level of primary sequence and expression control from syntenic loci, but also at the level of structure and function^1^. This leads to conservation of function despite divergent sequence and secondary structure among lncRNA orthologs^1^ and highlights the need to look beyond the sequence to understand the relationship between sequence, structure, and function in lncRNAs. The presence of modular secondary structures in several functional lncRNAs^2–4^ also supports the notion that lncRNA evolution favors the conservation of functionally relevant short stretches of sequence and/or structure that form distinct interacting domains for RNA–DNA, RNA–RNA, and RNA–protein regulatory complexes^5,6^. The combination of these different domains provides the lncRNAs with their unique functional specificity and diversity^5^. Such “modular RNA code” was validated in recent studies where short structure motifs were found to control the lncRNA stability and function, such as 3ʹ end triple helix in MALAT1 and NEAT1^7,8^, 5ʹ asymmetric G-rich internal loop (AGIL) motif in mouse Braveheart (Bvht)^4^, pseudoknots in human MEG3^9^, and a terminal hairpin stem-loop (SL1) in mouse lncRNA antisense to *Uchl1* mRNA^10–12^. In the antisense to *Uchl1*, the crucial SL1 region is within an embedded short interspersed nuclear element (SINE) from the B2 family, which forms the effector domain (ED) that is essential for targeted mRNA translation upregulation^10,11^.

SINEs are on average 100–300 bp in length and are non-autonomous retrotransposons that are evolutionarily derived from RNA polymerase III transcripts, mainly 5S ribosomal RNA (rRNA), 7SL RNA, and transfer RNA (tRNA)^13,14^. These elements, previously considered “junk DNA”, are abundant in gene-rich regions and their role in gene regulatory networks has been established^15–18^. Many SINEs are transcribed as enhancer RNAs and form the regulatory domains of functional lncRNAs^19–22^ as well as participating in the nuclear retention of lncRNAs^23^, STAU1-mediated mRNA decay^24^, and promotion of sense mRNA translation by antisense lncRNA^10^, thus supporting the Repeat Insertion Domains of LncRNAs (RIDL) hypothesis^6^. Yet the majority of SINE regulatory functions and structural features remain poorly understood. In mouse genome, the B2 family of SINE constitutes approximately 2.4% of the genome with presence of ∼3.5 x 10^5^ copies^25^. The SINEB2 family is broadly classified into B3A, B3, B2_Mm2, B2_mm1t, and B2_mm1a sub-families in descending order of evolutionary age according to the Repbase repeat elements database^26–28^. Synthetic antisense long non-coding RNAs SINEUPs which comprise a SINEB2 retrotransposon positively regulate the translation of the sense mRNA that they overlap^29,30^. Not only several mouse SINEB2s but also human free right Alu monomer (FRAM) repeat element and human MIRb are found in natural antisense lncRNAs and display post-transcriptional SINEUP-induced gene regulation^10,31^, despite their lack of similarity in terms of repeat length, sequence composition, and predicted secondary structure.

Here, to comprehend more clearly the extent of the SINEUP family, we used a synthetic SINEUP system to functionally screen 15 SINEB2 elements originally embedded in mouse natural antisense transcripts. The proven importance of SINE repeats in functional lncRNA domains prompted us to study the primary sequence and secondary structure of these SINE-derived RNAs. Although we have previously determined the folding of antisense *Uchl1* SINEB2 in vitro from its chemical footprinting and NMR data^11,12^, structural studies in solution cannot fully grasp the structural dynamics and interaction with cellular machinery. Moreover, to describe the relationship between sequence, structure, and function in SINE-derived RNAs, we need to study the RNA structure of multiple SINE-derived RNAs in the cellular environment and its specific compartments, because RNA can acquire different conformations depending on subcellular location, functional state, and biomolecular interactions^32^. Additionally, we need to understand RNA modifications in functional SINE-derived RNAs, specifically pseudouridylation, which is reported to be crucial for the structural stability, flexibility, and function of RNAs involved in translation^33–35^.

LncRNA structures are known to act as a molecular scaffold for proteins^36,37^, and in fact, the binding of heterogeneous nuclear ribonucleoprotein K (HNRNPK) with SINEUP and target mRNA complex is vital for SINEUP-mediated upregulation of target mRNA translation^38^. Previously, we found that upon HNRNPK knock-down, the co-localization of endogenous *Uchl1* mRNA and SINEUP RNA in the cytoplasm was decreased which resulted in loss of translation upregulation by SINEUP-UCHL1^38^. This negative effect of low HNRNPK expression on SINEUP function was also observed in SINEUP-GFP^38^. Furthermore, RNA immunoprecipitation analysis showed that HNRNPK binds with SINEUP-EGFP mRNA complex in both nucleus and cytoplasm and co-sediments with SINEUP and EGFP mRNA in free 40S, monosome and light polysome fractions in polysome profiling^38^. In other words, HNRNPK not only aids in the export of SINEUP and target mRNA complexes to the cytoplasm but also participates in SINEUP RNA-translation complex^38^, thus, exhibiting two discrete functions in the nucleus and the cytoplasm. This knowledge makes it imperative to examine how specific SINE RNA structures in different cellular compartments interact with RNA binding proteins (RBPs) and support their distinct functions. In addition to intramolecular structure, intermolecular interactions with other RNAs play a critical role in gene regulation by non-coding RNA^40,41^, notably the binding to rRNAs by linear and circular internal ribosome entry sites (IRESs) for promotion of translation^42,43^. In fact, our previous molecular dynamics simulation study on NMR solution structure of SINEB2 SL1 identified motifs in three-dimensions (3D) similar to those in rRNAs^44^. This data and the association of SINEUPs with ribosomal subunits in polysome profiles^38^ made us ponder the possibility that direct interaction between SINE-derived RNAs and rRNAs may be a mechanism by which SINEUPs enhance mRNA translation.

To solve the puzzle of SINE-transcribed RNA structure, we utilized a method called in vivo click selective 2ʹ-hydroxyl acylation and profiling experiment (icSHAPE), which chemically probes flexible RNA bases inside living cells and thus provides an advantage over other chemical or enzymatic methods^45^. We present two-dimensional (2D) secondary structure models for 17 mouse SINEB2s-derived RNAs and a human SINE FRAM RNA, providing for the first time experimentally verified in-cell 2D-structure models of evolutionarily close and distant SINE-transcribed RNAs at this scale. We found that structural deletion leading to the gain of an apical stem-loop (known as SL1) restores the activity in a SINEUP that is functionally dormant (i.e., lacking the ability to significantly enhance the target mRNA translation). Based on the 2D models, we discovered seven short conserved-structure motifs among the SINE RNAs tested, three of which harbor modified RNA bases. The icSHAPE-guided structures for the nucleus and cytoplasm showed that SINE RNAs in both mouse (SINEB2, derived from tRNA) and human (FRAM, derived from 7SL RNA) rearrange into a cloverleaf-like structure in the cytoplasm and have partially conserved structure motifs. We also observed that compartment-specific structures of SINE RNA are associated with distinctive binding patterns of HNRNPK in the nucleus and the cytoplasm by using a single-end enhanced crosslinking and immunoprecipitation assay (seCLIP)^46^. Additionally, when motifs were compared in silico with those in the Protein Data Bank (PDB), we detected a remarkable similarity of SINEUP 2D structure motifs with rRNA structure motifs. We also checked RNA–RNA pairs by the psoralen analysis of RNA interactions and structures (PARIS2) method^41^ and identified interactions of mouse and human SINE-derived RNAs with human 18S and 28S rRNAs in the conserved structure motif regions, which shed some light on SINEUP’s mechanism of action. Finally, we used our experimental 2D structure data to guide the prediction of structure models for 18 SINE-transcribed RNAs in 3D which can provide a base for future experimental 3D structure studies. In brief, this study demonstrates that despite sequence variability among SINE-transcribed RNAs, short stretches of conserved sequence and structure preserve the regulatory functions by providing binding sites for RBPs and rRNA. These short motifs are likely to be stabilized by modifications and non-canonical base interactions and are evolutionarily favored to maintain the SINEUP functionality even in distantly related SINE-derived RNAs.

## Results

### The SINEUP effect is a prevalent functional feature of mouse SINEB2s-derived RNAs

To experimentally verify the translation-regulatory function of various mouse SINE-transcribed RNAs in the SINEUP effector domain (ED), we used a previously established synthetic SINEUP-GFP model that enhances EGFP mRNA translation (**Fig. 1a**). We swapped the SINEB2 from antisense *Uchl1* in SINEUP-GFP with those from other mouse antisense to genes: *Txnip*, *Gadd45α*, and *Uxt* (**Supplementary Table 1a**). RepeatMasker revealed the presence of an inverted SINEB2 repeat spanning from 1389 to 1495 nucleotide positions (107 bp long) in antisense *Txnip*, and both antisense *Gadd45α* and antisense *Uxt* lncRNAs contain two inverted SINEB2 repeats in the region not complementary to the sense mRNA. Both antisense *Uxt* SINEB2 sequences are from the B3 sub-family; in the FANTOM3 cDNA clone, one sequence covered positions 160 to 290 (131 bp) and the other 774 to 960 (187 bp) (**Supplementary Table 1a**). Based on their natural order of occurrence from the 5ʹ end of the transcript, the sequences were named antisense *Uxt* SINEB2-a (131 bp, proximal to the 5ʹ end) and antisense *Uxt* SINEB2-b (187 bp, distal to the 5ʹ end) (**Supplementary Table 1a**, **Fig. 1b**). In contrast, both SINEB2 repeats of antisense *Gadd45α* belong to the B2_Mm2 sub-family (according to RepeatMasker nomenclature). We named these two sequences antisense *Gadd45α* SINEB2-a (131 bp, proximal to the 5ʹ end) and antisense *Gadd45α* SINEB2-b (114 bp, distal to the 5ʹ end) (**Supplementary Table 1a**). To understand whether both SINEB2 sequences are essential for SINEUP function and how their order affects SINEUP activity, we designed antisense *Gadd45α* and antisense *Uxt* constructs with a or b sequences and also with both sequences in the natural (ab) or the inverted (ba) order (**Fig. 1b**). The shortest SINEB2 of those tested, antisense *Txnip* SINEB2 (107 bp), enhanced the GFP protein level (**Fig. 1c** and **d**, **Supplementary Table 1**). None of the four constructs with antisense *Gadd45α* SINEB2s could exert the SINEUP effect, regardless of the order of SINEB2 a and b (**Fig. 1c** and **d**). Indeed, natural antisense *Gadd45α* was found to inhibit translation of sense *Gadd45α* (**Extended Data Fig. 1**), which implies that the SINEUP activity of antisense *Gadd45α* SINEB2 RNA is dormant, possibly due to the lack of functional regions. Interestingly, of the antisense *Uxt* SINEB2 repeats, only SINEB2-b upregulated the GFP protein level significantly, and when the position of the two SINEB2 sequences was switched from the natural order (ab switched to ba), their SINEUP effect was obliterated (**Fig. 1c** and **d**). As negative controls, in addition to empty vector, we tested direct antisense *Uchl1* SINEB2, where the orientation of SINEB2 was flipped from the one in the functionally active construct, and the deletion mutant of antisense *Uchl1* SINEB2 (ΔAS *Uchl1* SINEB2) in which the deletion of SINEB2 shifts downstream synthetic and non-SINEB2 annotated sequences in its place in the ED of SINEUP^10^. Consistent with our previous work^10^, none of these negative controls could upregulate GFP translation which demonstrates that SINE RNA region is essential for SINEUP function (**Fig. 1c** and **d**). Upholding the post-transcriptional regulatory nature of SINEUPs, none of the SINEUPs changed the EGFP mRNA expression level significantly (**Extended Data Fig. 2a**).

**Figure 1.**
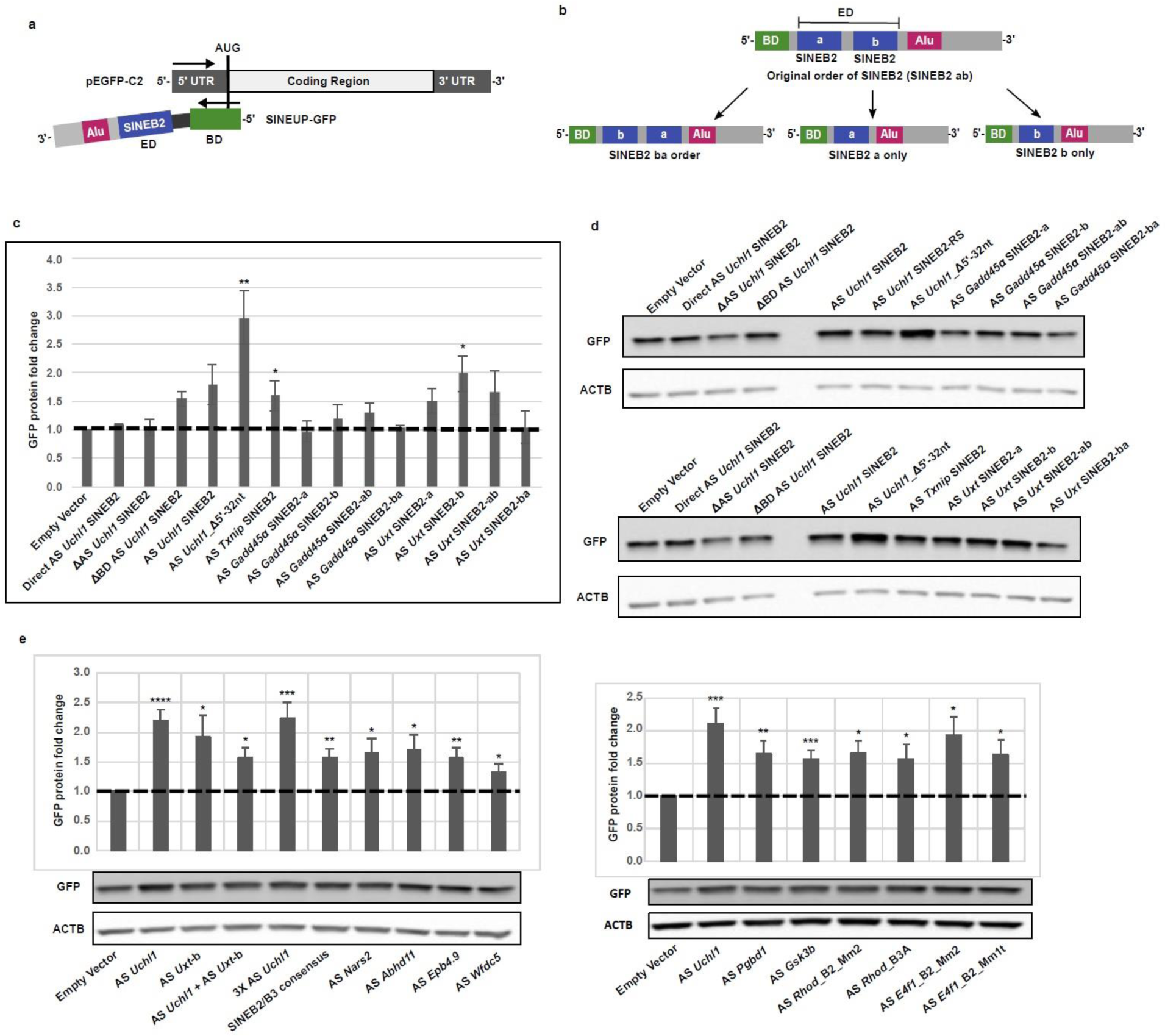
SINEB2 RNAs from different mouse antisense lncRNAs can act as SINEUP effector domain and upregulate GFP protein expression in HEK293T/17. (a) Schematic representation of the sense/antisense EGFP system used to validate the effect of different SINEB2 RNAs on translation. The SINEUP binding domain (BD) is shown in green. The inverted SINEB2 region is the effector domain (ED) shown in blue. Alu repeat and 3ʹ tail of SINEUP are shown in magenta and gray respectively. (b) SINEUP construct design to verify independent and combined effects of two SINEB2 elements from the same antisense lncRNA. (c) Western blot–based EGFP band intensities normalized to ACTB expression as a measurement of SINEUP-induced protein fold change observed in the HEK293T/17 cell line at 24 h post-transfection. Empty vector, Direct AS Uchl1 SINEB2, and ΔAS Uchl1 SINEB2 (SINEB2 deletion) wereused as controls. (d) Western blot images with anti-GFP and anti-ACTB antibodies. ACTB was used as a loading control. Abbreviations: AS, antisense; Direct, direct orientation of SINEB2; Δ, deletion; a and b, two different SINEB2 elements from the same antisense lncRNA; ab and ba, the two SINEB2 elements in original and reversed order, respectively; AS Uchl1 SINEB2–RS, antisense *Uchl1* SINEB2 construct without any restriction enzyme sites around the SINEB2 elements. (e) GFP protein fold change in HEK293T cells after co-transfection with sense EGFP and miniSINEUP (at 24 h post-transfection). Western blot images and corresponding GFP band intensities (normalized to ACTB expression level) calculated using ImageJ software. An empty vector was used as the negative control and antisense *Uchl1* SINEB2 containing miniSINEUP (shown as AS *Uchl1*) as the positive control. Sample labels indicate names of antisense lncRNAs from which the SINEB2 sequences were isolated, which represent four sub-families (based on RepeatMasker annotation): B3, B3A, Mm2, and Mm1t. SINEB2 elements are from the B3 sub-family unless specifically stated. AS *Uchl1* + AS *Uxt*-b is the combination of AS *Uchl1* SINEB2 and the second SINEB2 of AS *Uxt*; 3× indicates three repeats of SINEB2; and SINEB2/B3 consensus is the B3 sub-family consensus sequence taken from the RepBase database. N = 3 (for c) and n = 5 (for e); error bars ± standard error mean; **** P < 0.00005; *** P < 0.0005; ** P < 0.005; * P < 0.05 by two-tailed Student’s *t*-test.

To further expand analysis of the SINEB2-derived RNAs present in mouse natural antisense transcripts (**Supplementary Table 1**), we used the shortest functional model of SINEUP-GFP, named miniSINEUP-GFP^29^. SINEB2 sequences from antisense *Uchl1* and antisense *Uxt* enhanced GFP protein expression by around 1.8–2.2-fold (**Fig. 1e**, Lanes 2 and 3 in the left graph). We additionally found that combining the SINEB2 from antisense *Uchl1* with antisense *Uxt* SINEB2-b upregulated GFP protein production; however, the protein fold change was lower than the effect of either SINEB2 independently (**Fig. 1e**, Lane 4 in the left graph). We additionally tested three repeats of antisense *Uchl1* SINEB2 but the SINEUP function remained similar to that with only one repeat, implying that these repeats do not function in a synergistic or additive manner (**Fig. 1e**, Lane 5 in the left graph).

Since the SINEB2 sub-family B3 acted as a functionally effective SINEUP ED, we decided to check the SINEUP potential of consensus subfamily B3 sequence from the Repbase database^27,47^. Indeed, the sequence is functionally active against GFP (∼1.6-fold change) (**Fig. 1e**, Lane 6 in the left graph), but not to the level of antisense *Uchl1* SINEB2 RNA, which is a similar result to tests on other SINEB2 sequences in the B3 sub-family from antisense lncRNAs to genes: *Nars2*, *Abhd11*, *Epb4.9*, *Wfdc5*, *Pgbd1*, and *Gsk3b* (**Fig. 1e**, Lanes 7–10 in the left graph and Lanes 3–4 in the right graph).

The failure of SINEUPs with antisense *Gadd45α* SINEB2 from the B2_Mm2 sub-family prompted us to test whether the SINEUP effect is limited to the B3 sub-family only. To confirm this, we investigated two B2_Mm2 sub-family sequences (from antisense *Rhod* and antisense *E4f1*), one B2_Mm1t (from antisense *E4f1*) and one B3A (from antisense *Rhod*) (**Fig. 1e**, Lanes 5–8 in the right graph) and found that all the SINEB2 sequences, regardless of their sub-family type, length, or parent natural antisense transcripts, upregulated GFP protein expression to more or less the same extent, which reveals the vastness of the SINEUP family. Similar to that of long SINEUPs, this miniSINEUP effect was also manifested at the post-transcription level (**Extended Data Fig. 2b**). Although the SINEUP RNA level for SINEB2s from antisense *Wfdc5*, *Pgbd1*, and *Gsk3b* was unusually high (3–10-fold higher than other miniSINEUPs), the GFP mRNA expression level in all miniSINEUPs remained unaltered.

### Functional SINEs form double-stranded structures in the cell and consist of conserved short structure motifs

To check any common sequence features between different SINEB2-transcribed RNAs showing the SINEUP effect and to identify any functionally important regions, we performed multiple sequence alignment using ClustalW (**Extended Data Fig. 3a** and b). This revealed conserved regions of RNA pol III internal promoters known as A box and B box^48^ (**Extended Data Fig. 3a**) along with sequence diversity between SINEB2 elements tested in this study. Moreover, in the phylogenetic tree based on the multiple sequence alignment, the functionally active SINEB2s that significantly upregulated GFP translation were not clustered separately from the inactive SINEB2s that failed to do so (**Extended Data Fig. 3b**). In other words, on the basis of sequence, the functional SINEB2 RNAs do not possess any unique features that confer a functional advantage over the inactive SINEB2 RNAs. This lack of conserved sequence domains led us to examine the in-cell secondary structure of SINEB2s by using the icSHAPE method to map the RNA secondary structure inside living cells^45,49^.

We compared the icSHAPE antisense *Uchl1* SINEB2 RNA secondary structure with our previous in vitro chemically probed structure^11^, which had identified four internal-loops (IL1–IL4) and three stem-loops (SL1–SL3) (**Fig. 2a**). Both methods gave a notably similar double-stranded structure for antisense *Uchl1* SINEB2 RNA (**Fig. 2a**). The icSHAPE-based structure comprises three internal loops similar to the IL2, IL3, and IL4 found in the chemical footprinting-based structure, with slight differences in the sequence composition. The in-cell structure lacks SL3, which indicates the influence of intracellular factors and dynamic structural changes compared to the less complex conditions in vitro. Interestingly, SL1 and SL2 are conserved between the structures derived by these two different methods, which suggests that they represent the most dominant and stable structure components in the ensemble. The structural conservation is also understandable knowing that SL1 is crucial for SINEUP function^11^. In a similar way, secondary structure modelling was performed for other SINEB2s-derived RNAs (**Fig. 2b** and **c**, **Extended Data Fig. 4**) and revealed that they all retained double-stranded features with a terminal GC-rich stem-loop similar to SL1 of antisense *Uchl1* SINEB2, even though all SINEB2 RNAs tested here had diverse global structures with variable sequence length and composition.

**Figure 2.**
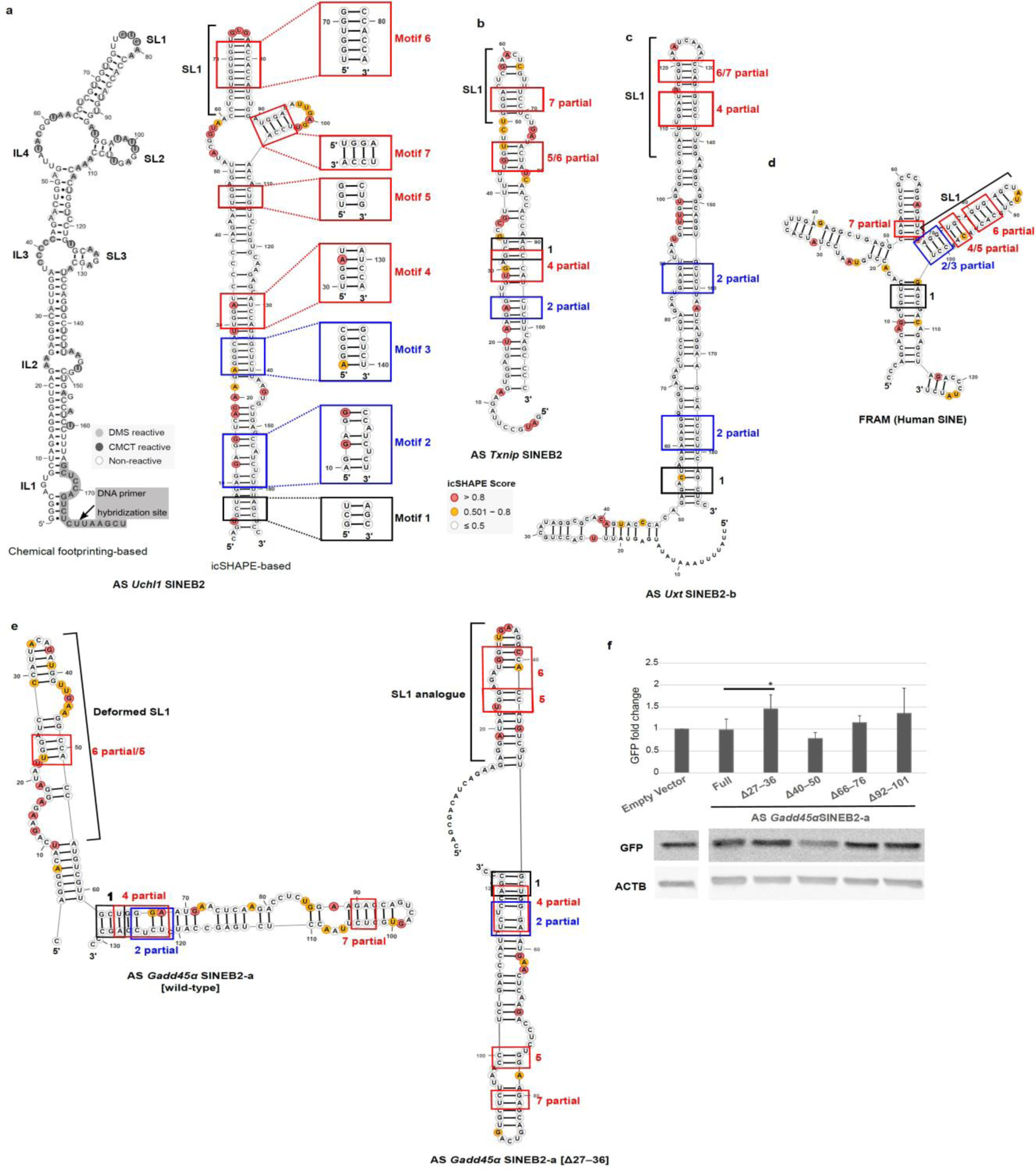
In-cell SINE structures and conserved motifs in functionally active SINEUPs. (**a**) Antisense (AS) *Uchl1* SINEB2 RNA secondary structure based on icSHAPE data (right) is compared with the one previously identified using chemical footprinting^11^. In chemical footprinting-based structure, DNA primer hybridization site is highlighted in gray and indicated by an arrow; light and dark gray circles represent DMS and CMCT reactive bases, respectively; non-reactive bases are outlined by a circle. In icSHAPE-based structure, squares highlight the common sequences and structure motifs in in-cell structures of SINE, with their corresponding motif number. AGG (blue) and UGG (red) motifs are shown. (**b**–**d**) Location of SINEUP motifs in icSHAPE-derived structures of functionally active SINE RNAs from (**b**) antisense *Txnip* SINEB2, (**c**) antisense *Uxt* SINEB2-b and, (**d**) human SINE element FRAM. (**e**) icSHAPE-derived RNA secondary structures and local SINEUP motifs (colors as in Figure 2a) of functionally dormant wild-type antisense *Gadd45α* SINEB2-a and its deletion mutant antisense *Gadd45α* SINEB2-a [Δ27–36] with gained SL1. (**f**) Effect of antisense *Gadd45α* SINEB2-a deletion mutants on GFP protein expression as measured by Western blot. An empty vector was used as the negative control. Four mutants with random deletion, as listed in the Figure, were compared with the full-length construct (Full). N = 3, error bars ± standard deviation, * P < 0.05 by two-tailed Student’s *t*-test. AS, antisense; Δ, deletion; SL1, stem-loop 1. Nucleotide color indicates normalized icSHAPE reactivity score: red, ≥ 0.8; yellow, from above 0.5 to 0.8; white encircled with grey, ≤ 0.5; bases not encircled, no icSHAPE data available.

The importance of short structure motifs like SL1 in the SINEUP function prompted us to look for other local motifs that are shared between different SINEB2-transcribed RNAs. For this purpose, we utilized the ExpaRNA tool^50^ and, using the antisense *Uchl1* SINEB2 RNA structure as a base, searched for its exact sequence and structure patterns in other icSHAPE-derived SINEB2 RNA structures. From this search, we discovered seven GC-rich structure motifs partially conserved among the SINEB2 RNA structures tested in this study (**Fig. 2a**–**c**, **Extended Data Fig. 4**). Motifs 2 and 3 are comprised of ‘AGG’ trinucleotide(s), while motifs 4, 5, 6, and 7 are ‘UGG’ enriched. Motif 6 lies within the SL1 region making it a potentially functional motif. Furthermore, we wondered if the motifs are conserved in SINE-transcribed RNAs that are evolutionarily distant from mouse SINEB2 elements, such as human SINE FRAM, which has been proven to be effective as a SINEUP against both endogenous and exogenous mRNA targets^31^. Indeed, the structure motifs in the whole-cell icSHAPE structure of FRAM RNA provided five matching regions (**Fig. 2d**). This hints that conservation of such short motifs may contribute in functional preservation. We further examined the PDB database for 3D RNA structure motifs that were akin to our SINEUP structure motifs. For this purpose, we employed a sequence- and structure-specific search for each of the seven SINEUP motifs in the RNA Characterization of Secondary Structure Motifs (CoSSMos) database^51^. This resulted in hits that matched 16S rRNA (PDB ID: 5JB3) and 18S rRNA (PDB ID: 5A2Q) to 2D structures for SINEUP motifs 2 and 6, respectively (**Extended Data Fig. 5a**). Interestingly, the matching motifs are very similar to in-cell derived SINEUP motifs in terms of sequence and structure. On closer inspection, we observed that the matching region to 18S rRNA is within helix 34, which forms the head of the mRNA entry channel^52^ (**Extended Data Fig. 5b**).

### Acquisition of SL1-like structure turns a dormant SINEUP active

Considering the conservation of short local structure motifs in functional SINE-transcribed RNAs, we theorized that dormant SINEUPs, here the SINEB2s that failed to significantly upregulate GFP protein expression in **Fig. 1c**, might include embedded inhibitory regions that prevent their proper folding into functional domains. To test this, we chose a functionally inactive SINEB2 sequence, antisense *GADD45α* SINEB2-a, and checked the effect of deletion of different sequence regions on its structure. In total, we created four deletion mutants at positions 27–36, 40–50, 66–76, and 92–101 from the 5ʹ end of SINEB2 including two tRNA-derived conserved internal RNA polymerase III promoter regions, A-box and B-box (**Extended Data Fig. 3a**). We observed structural changes caused by these deletions by in-silico RNA structure prediction which we later confirmed in the cell by icSHAPE. Analysis of in-cell structures of deletion mutants in comparison to that of wild-type antisense *GADD45α* SINEB2-a and antisense *Uchl1* SINEB2 (**Fig. 2a, e**, **Extended Data Fig. 6**) showed that out of all the mutants, Δ27–36 shares the most sequence and structural similarity in SL1 structure and motif 6 with functional antisense *Uchl1* SINEB2 (**Fig. 2a, e**). The wild-type antisense *Gadd45α* SINEB2-a carries a deformed SL1 where the stem is interrupted by multiple internal loops (**Fig. 2e**). Upon deletion of 27–36 nucleotide positions, this stem becomes less structured and transforms into a properly folded SL1 similar to antisense *Uchl1* SINEB2. The Δ40–50 mutant, though shares similarity in overall 2D structure in terms of position of motifs 1, 2, 4, 5, and 7 with Δ27–36 mutant, lacks a proper SL1 and no motif 6 (**Extended Data Fig. 6**. On the other hand, Δ66–76 and Δ92–101 mutants contain partial motif 6 but have truncated SL1 (**Extended Data Fig. 6**). This disparity in structure is reflected in their SINEUP function. When we co-transfected the mutants with sense EGFP into HEK293T/17 cells, as done in the previous SINEUP-GFP functional analysis experiment we observed that out of the five mutants Δ27–36 could significantly upregulate GFP protein level compared to full-length antisense *GADD45α* SINEB2-a (**Fig. 2f**). Deletion mutant Δ40–50 and Δ66–76 could not upregulate GFP translation while Δ92–101 mutant in some experiments showed around 1.5-fold increase in GFP protein but had inconsistent performance in others (**Fig. 2f**). This suggested that some structural features in the wild-type antisense *GADD45α* SINEB2-a are detrimental to function, whereas the acquisition of short local structure motifs leads to functionally favorable RNA folding, which can restore SINEUP function. Especially, properly folded SL1 and motif 6 are essential not only for exerting the SINEUP effect but also for ensuring functional stability.

### SINEUP structure dynamics across nucleus and cytoplasm

Our SINEUP icSHAPE data from whole cell lysate showed a set of common motifs among different SINEB2s, yet it was an average of whole-cell structure ensemble and did not help to infer structure dynamics within the cell. RNA structure is closely related to cellular processes and can acquire distinct conformations depending on the cellular compartment^32^. Understanding the different RNA structures becomes more crucial in the case of SINEUPs because they are involved in post-transcriptional translation regulation, a function specific to cytoplasm. Therefore, it is logical to examine the structure dynamics of functional SINEB2s-derived RNAs both in the nucleus and the cytoplasm, to better understand the contribution of structural conformations to the biological processes specific to these subcellular regions. To achieve this, we made icSHAPE libraries from nucleoplasmic and cytoplasmic fractions of cells transfected with SINEUP-GFP carrying either mouse antisense *Uchl1* SINEB2 or human SINE RNA FRAM. The 2D structure models based on this icSHAPE data revealed that the base of the SINEB2 RNA structure is mostly stable between the nucleus and the cytoplasm, with only a slight variation observed around the region of motif 2, being slightly more open in the nucleus than in the cytoplasm (**Fig. 3a, b**), and a similar structure difference in human FRAM-transcribed nuclear RNA (**Fig. 3c, d**). In addition, the internal loop between motifs 5, 6, and 7 in the nuclear mouse SINEB2 RNA goes through a structural rearrangement in the cytoplasm and transforms into a cloverleaf-like structure (**Fig. 3a, b**). In the case of human FRAM RNA, both nuclear and cytoplasmic structures are quite similar and form a cloverleaf-like structure, though the cytoplasmic structure is slightly more structured (**Fig. 3c, d**). This could be possibly due to the transient nature of any alternative structures, as icSHAPE structure represent only the average of the structure ensemble.

**Figure 3.**
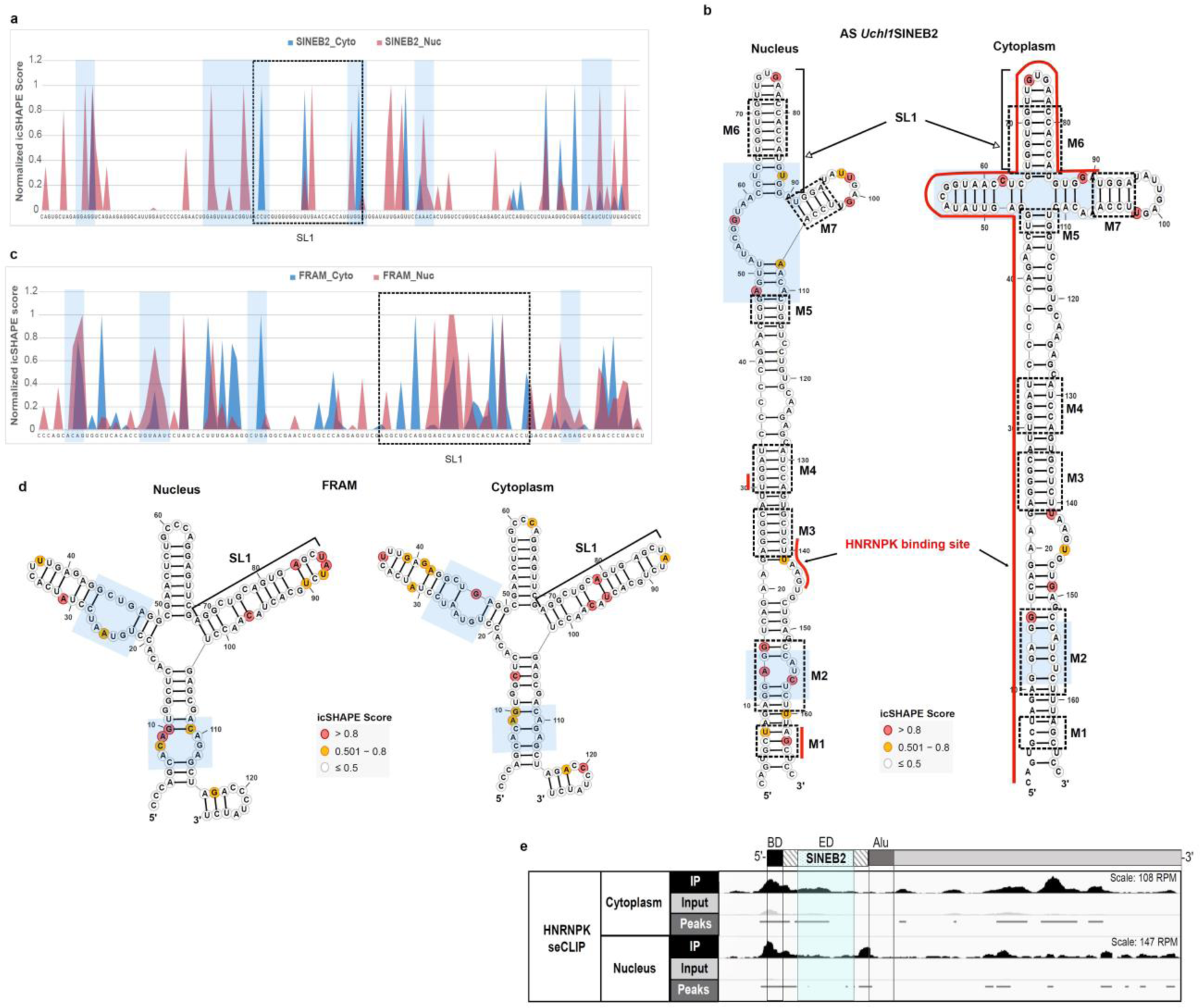
SINEUP structure dynamics and interaction with HNRNPK in the nucleus and cytoplasm. (**a**) Normalized icSHAPE reactivity score profiles of mouse antisense (AS) *Uchl1* SINEB2 RNA in the nucleus and cytoplasm and (**b**) related 2D structure models created from icSHAPE data. SINEUP motifs from M1 to M7 are indicated by dashed squares. (**c**) Normalized icSHAPE reactivity score profiles of human SINE RNA FRAM in the nucleus and cytoplasm and (**d**) their respective 2D structure models. Different structural regions between the compartments are shaded in blue boxes. (**e**) seCLIP data for HNRNPK binding to SINEUP-GFP RNA (of AS *Uchl1* SINEB2) in the nuclear and cytoplasmic fractions of HEK293T cells. IP: immunoprecipitation against HNRNPK; input: not immunoprecipitated, contains all RNA-protein complexes in the given fraction. Normalized peak regions representing significant enrichment of HNRNPK binding site in IP compared to input are indicated as horizontal bars. The SINEB2 region is highlighted in blue. BD, binding domain; ED, effector domain. Identified HNRNPK binding sites are marked by a red line on 2D structure models of AS *Uchl1* SINEB2 in panel b.

The nucleus-specific internal loop structure might play a role in the export of SINEUP RNA to the cytoplasm, whereas the cytoplasmic cloverleaf-like structure with SL1 might be crucial for interaction with translation machinery. Interaction with HNRNPK is crucial in both of these SINEUP processes^38^. To check how the different nuclear and cytoplasmic structures of SINEB2 RNA are associated with HNRNPK, we prepared seCLIP^46^ libraries from the nuclear and cytoplasmic fractions of SINEUP-GFP co-transfected cells (**Fig. 3e**). The results showed that in the nucleus HNRNPK interacts with short stretches of SINEB2 coinciding in motifs 1, 3, and 4 (**Fig. 3b**), but in the cytoplasm HNRNPK occupied a long double-stranded region covering the 5ʹ end of the stem in all the motifs except motif 7. Such disparate interactions could possibly arise from distinct functions of HNRNPK in SINEUP RNA transport and in translation that are specific to the cellular compartment.

### SINEUP structure motifs interact with rRNA

We wanted to further investigate the role of SINEB2 RNA structure in translation because it has features reminiscent of tRNA; it forms a cloverleaf-like structure in the cytoplasm and possesses two pseudouridine (Ψ) sites at 16 bp and 112 bp (**Extended Data Fig. 7**). CircRNA IRESs^43^ and many other IRESs are known to interact with 18S rRNA^42,53^ to induce cap-independent translation, however the region by which SINEUP-RNA interacts with ribosomal subunits and polysomes is not known.

Considering the similarities in sequence and structure, we wondered whether such motifs in SINEB2 RNA can interact with rRNA. To get a better understanding of SINEUP–rRNA interactions, we employed PARIS2, which has proved useful in capturing direct mRNA–rRNA short interactions^41,54^. We prepared PARIS2 libraries from HEK293T/17 cells transiently expressing SINEUP-GFP (antisense *Uchl1* SINEB2) or miniSINEUP-GFP (FRAM) along with their target sense EGFP mRNA and sequenced them on a next-generation sequencing platform. We could capture few incidents of SINEUP-rRNA pairs (**Fig. 4**, **Fig. 5**, **Extended Data Fig. 8**), which is understandable given the technical limitation of psoralen crosslinking and inefficient detection of scarcely expressed RNA duplexes and highly dynamic interactions in PARIS2^41,55^. Nevertheless, it was intriguing to see interactions of mouse SINEB2 element with human rRNA in both small and large subunits of ribosome (**Fig. 4a, b**, and **c**). We discovered that the 3ʹ side of motifs 1 and 2 of SINEB2 RNA interacted with a region in helices 14 and 15 of 18S rRNA (Int 1 in **Fig. 4a** and **c**). On the other hand, the region between motifs 5 and 6 formed contacts with domain III helices 55, 56, and 59 of 28S rRNA. Motif 5 and the flanking region also interacted with domain I, helix 25, expansion segments ES7c and ES7d of 28S rRNA (Int 2 and 3 in **Fig. 4b** and **c**).

**Figure 4.**
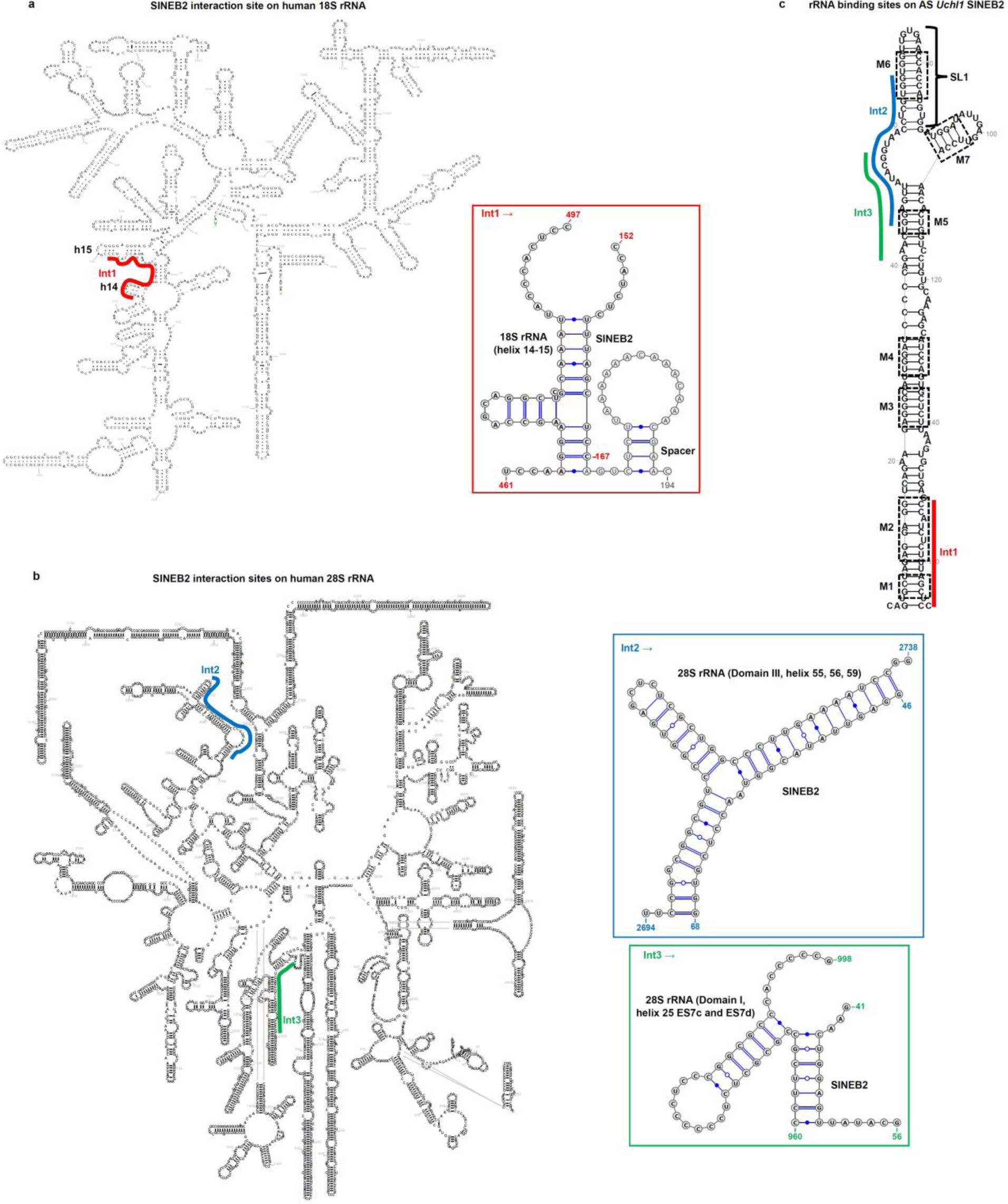
Mouse SINEB2 RNA interacts with human rRNA in cell. Antisense (AS) *Uchl1* SINEB2 interacting regions (Int) in human (**a**) 18S rRNA (Int1) and (**b**) 28S rRNA (Int2 and 3) identified by PARIS2. Different interaction sites are color-coded, and rRNA-SINEB2 pairs are shown in the corresponding color. Possible base-pairing based on sequence complementarity of interacting regions is shown. (**c**) SINEB2 regions interacting with 18S and 28S rRNA regions marked in (a) and (b) on icSHAPE data-driven SINEB2 structure from whole-cell lysate. SINEUP motifs from M1 to M7 are indicated by dashed squares.

**Figure 5.**
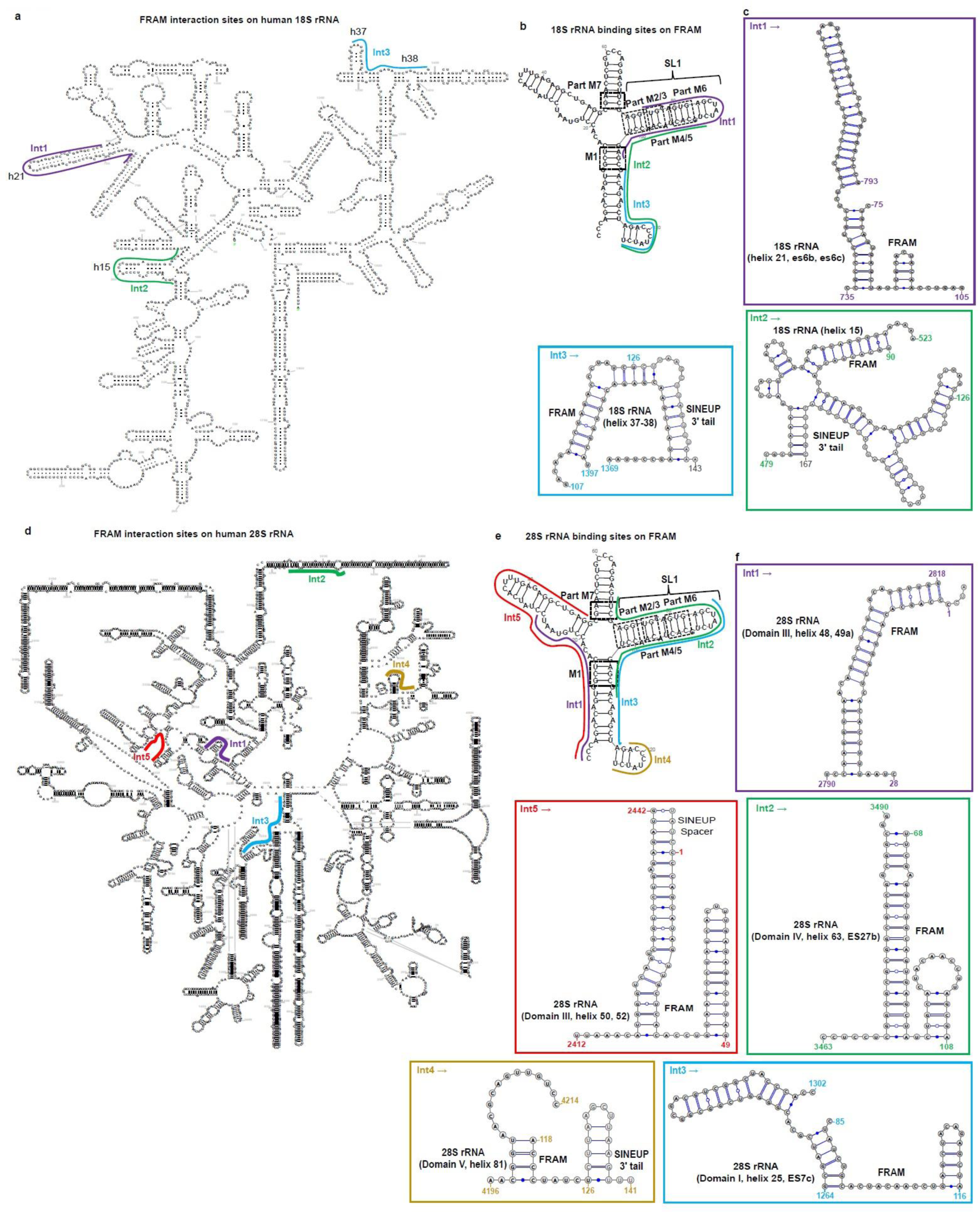
In-cell interaction of functional human SINE element FRAM with rRNA. (**a**) Regions of 18S rRNA found to pair with the SINEUP FRAM region in PARIS2 data. (**b**) Corresponding binding regions to (a) are highlighted on FRAM RNA. (**c**) Description of FRAM-18S rRNA pairs and interaction sites marked in (a) and (b). (**d**) 28S rRNA regions identified as FRAM interaction sites by PARIS2 and (**e**) their corresponding binding regions on FRAM. (**f**) Details of FRAM-28S rRNA interactions highlighted in (d) and (e). Different interaction sites (Int) are represented by different colors. Possible base-pairing based on sequence complementarity of RNA-RNA pairs is shown in (c) and (f). In (b) and (e) the icSHAPE-guided FRAM structure from whole cell lysate is shown. Matching regions to SINEUP motifs from M1 to M7 are indicated by dashed squares.

Interestingly, functionally active human SINE-derived RNA FRAM displayed conservation of this rRNA interaction trait. With better expression of FRAM RNA, we were able to detect more instances of FRAM-rRNA interaction than for SINEB2 RNA. Similar to SINEB2, the 3ʹ side of motifs 1 and 2 and of motifs 5 and 6 of FRAM RNA formed contacts with helix 15 of 18S rRNA (Int 2 in **Fig. 5a, b, c**). In addition, the FRAM RNA stem loop analogous to SL1 (containing motifs 2/3, 5, and 6) interacted with 18S rRNA helix 21, es6b and es6c (Int 1 in **Fig. 5a, b, c**), and helices 37 and 38 paired with the 3ʹ end region of FRAM RNA (Int 3 in **Fig. 5a, b, c**).

Moreover, the 5ʹ side of FRAM RNA flanking motif 1 interacted with 28S rRNA domain III helix 48 and 49a (Int1 in **Fig. 5d, e, f**), and helix 50 and 52 (Int5 **Fig. 5d, e, f**), whereas FRAM SL1 region interacted with 28S rRNA domain IV, helix 63, ES27b and with domain I, helix 25, ES7c (Int 2 and 3 in **Fig. 5d, e, f**). The short stem-loop at the 3ʹ end of FRAM structure was involved in interaction with 28S rRNA domain V, helix 81 (Int4 in **Fig. 5d, e, f**).

The presence of SINEUP-rRNA pairs in PARIS2 data in addition to our knowledge of SINEUPs’ fate in the polysomes^38^, that over 85% of the cytoplasmic SINEUP-GFP co-sediments with EGFP mRNA and HNRNPK in free/40S ribosomal subunit and monosome fractions, supports the notion of SINEUP-rRNA interaction^38^.

### SINE RNA 3D structure models reveal non-canonical intramolecular interactions

The SINE 2D structure models helped us to understand the conservation and functional importance of local short motifs in maintaining the SINEUP effect by interacting with RBP and rRNA. However, they lacked information about non-canonical intramolecular interactions that stabilize the RNA structure and affect the availability of interaction sites in 3D. This results in canonically unpaired regions in the 2D structures that are actually non-canonically paired in 3D^56^. Therefore, we took advantage of in vivo SINE RNA 2D structure information and incorporated it into the machine translation of 3D structure models. Thus, we created 3D models for 17 mouse and one human SINE-derived RNAs (**Fig. 6** and **Extended Data Fig. 9**). The 3D models revealed that the SINEUP motifs are stabilized by base-stacking interactions along the double-helical structure and that bases in the internal loop and stem-loop regions form non-canonical interactions adding more complexity to the structure (**Fig. 6 a,c,e,g** and **Supplementary Tables 2 and 3**). The molecular surface view of the nuclear SINEB2 RNA shows an elongated helical structure where SL1 takes the apical position (**Fig. 6b**). Interestingly, human nuclear FRAM RNA, which has a cloverleaf-like conformation in 2D (**Fig. 3d**), folds similarly to nuclear SINEB2 RNA in 3D, with the three arms positioned in such a way that all the motifs are in the central arm, SL1 is at the apex, and the other two arms are close to the central one (**Fig. 2d** and **Fig. 6 e,f**). In the cytoplasm, both SINEB2 and FRAM RNAs fold into a cloverleaf-like conformation where SL1 moves from the center to the left position and in FRAM RNA the other two arms open up (**Fig. 6 c,d,g,h**). We also analyzed the 3D conformation of functionally inactive antisense *Gadd45α* SINEB2-a and its deletion mutants in comparison to antisense *Uchl1* SINEB2 (**Extended Data Fig. 10**). The 3D models showed that among all the mutants, the functional Δ27–36 mutant adopts a similar global structure as well as SL1 and motif 6 sub-structures with that of antisense *Uchl1* SINEB2 in 3D space (**Extended Data Fig. 10a,c**) consistent with the 2D structure data. The wild-type antisense *Gadd45α* SINEB2-a and functionally inactive mutants Δ40–50, Δ66–76 and Δ92–101 which have deformed SL1, vary in RNA 3D architecture from the functional mutant (**Extended Data Fig. 10 a,b,d–f**).

**Figure 6.**
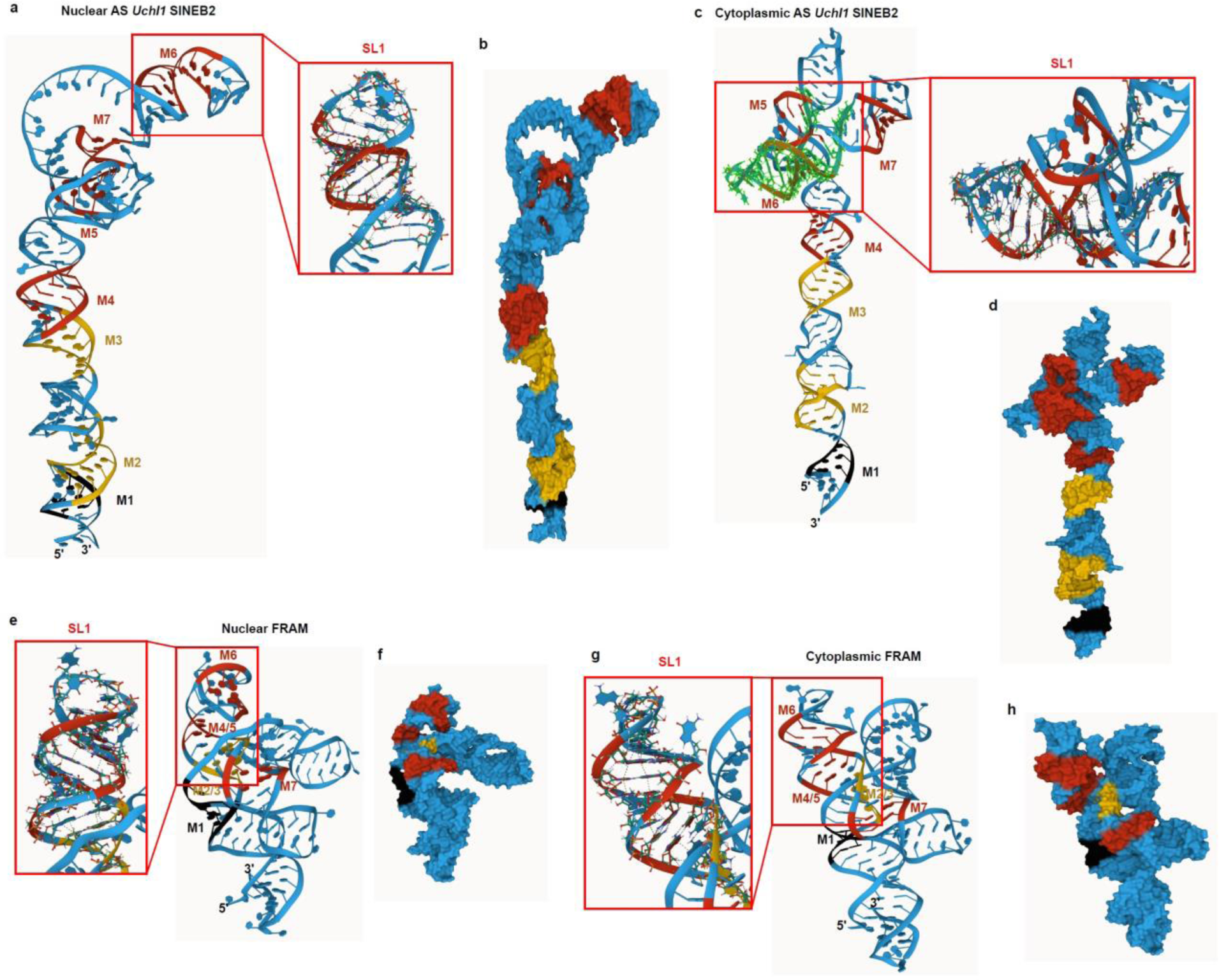
Machine-translated 3D RNA structure models based on in vivo SINE RNA 2D structures. 3D structure models for (**a**–**d**) mouse antisense (AS) *Uchl1* SINEB2 RNA in the nucleus (**a**,**b**) and cytoplasm (**c**,**d**) and for (**e**–**h**) FRAM RNA in the nucleus (**e**,**f**) and cytoplasm (**g**,**h**). SINEUP structure motifs M1–M7 are shown. Red frames show enlarged views of the SL1 region. Panels b, d, f, and h show the molecular surface view of the 3D models.

## Discussion

There are about 175 families of SINE scattered across different domains of life^57^. Although generally considered as parasite elements of the genome, a growing number of studies point to their gene-regulatory activity, such as enhancer-like function, polyadenylation signal, transcription factor binding sites, mRNA nuclear retention, and translation upregulation^10,15–18,23^. However, there are no extensive studies on the sequence and structural features of functional SINE-transcribed RNA elements. Here, we examined the SINEUP activity of 15 as-yet-uncharacterized SINEB2-derived RNAs from mouse natural antisense transcripts and assayed their function in GFP protein upregulation. Most (12 out of 15) of the SINEB2 RNAs showed SINEUP activity, but the remaining 3 (antisense *Gadd45α*-SINEB2-a and -b, and antisense *Uxt* SINEB2-a) failed to upregulate GFP protein expression significantly under the current experimental settings. The shortest SINEB2 RNA to enhance GFP protein expression, antisense *Txnip* SINEB2, was found to be 107 nucleotides long, which is half the length of the longest functional natural SINEB2-derived RNAs tested here (antisense *Nars2* SINEB2 and antisense *Epb4.9* SINEB2). This suggests that the SINEUP class of RNAs in nature is vast and diverse and is likely to be a much larger class of antisense RNAs than yet explored. Functional observation of the consensus SINEB2 sub-family B3 sequence also supported this, although it enhanced GFP translation to a lesser extent than the highly functional antisense *Uchl1* SINEB2 RNA. This might be because consensus sequences represent the most frequent occurrence of a particular nucleotide in a multiple sequence alignment of related sequences, which may not necessarily be functionally optimal. We observed that in most cases only one repeat of SINEB2 RNA as the effector domain is sufficient to exert the SINEUP effect, possibly because two SINEB2-derived RNAs (which are generally GC-rich) may share some short complementary regions that can interact and hinder the proper RNA folding into key functional features or may compete for binding proteins and RNA partners. Another interpretation of this non-synergistic nature of these functional SINEB2 RNAs is that SINEUP-induced protein upregulation is used to fine-tune translation in the cell, keeping the protein within a physiological range of up to 1.5–3-fold change. As previously observed, this fold change is sufficient to rescue haploinsufficient gene dosage and disease phenotype in vivo by upregulation of protein expression of the subunit 7B of cytochrome c oxidase (*cox*7b) in a medaka fish model of human microphthalmia with linear skin defects (MLS) syndrome^58^, and rescue of endogenous defective Frataxin (FXN) expression in a cellular model of Friedreich’s ataxia^59^ as well as rescue of neurodegeneration and motor phenotype by enhancing endogenous Glial cell-derived neurotrophic factor (GDNF) protein expression in a mice Parkinson’s disease model^60^. Incidentally, this characteristic makes SINEUPs ideal for therapeutic studies where proteins need to be regulated within a physiological range. The sequence diversity within the same B3 sub-family or in comparison with other sub-families such as B3A, B2-Mm2, and B2_Mm1t (as per Repbase nomenclature)^27^ does not prevent the SINEUP function. This observation confirms the universality of the SINEUP function in the SINEB2 repeat family and the flexibility of its members to maintain key functional features despite varying length and composition of RNA sequences. It would be interesting to explore other families of SINE elements in other species and across a wide variety of mammalian and eukaryotic genomes in the future.

Noting the sequence diversity of functional SINEB2-derived RNAs and the importance of RNA structure in non-coding RNA functions, we looked at the role of the SINE RNA secondary structure in the SINEUP phenomena in living cells by using icSHAPE experiments. Consequently, we noticed a high correlation between the in-cell 2D structure of antisense *Uchl1* SINEB2 RNA and the previously reported structure from chemical footprinting^11^. Moreover, we found that the key functional SL1 structure was conserved among various SINEB2s RNAs. When the functionally dormant antisense *Gadd45α* SINEB2-a RNA that lacks an SL1 region was mutated to change its structure, one of the mutants (Δ27–36) acquired a properly folded SL1 structure and a 2D structure similar to the icSHAPE structure of antisense *Uchl1* SINEB2 RNA, resulting in SINEUP activity gain. This further emphasizes the functional significance of this short structure in SINEUP-induced gene regulation.

We discovered 7 conserved structure motifs among the icSHAPE data-driven 2D structure models of all the SINEB2-derived RNAs tested here, though only partial matches in sequence were observed, given the sequence variation. Since we observed partial matches to these structure motifs in the functionally inactive SINEs also, we hypothesize that the SINEUP function is influenced not just by the mere presence of these structure motifs but also by their relative positions on the structure. Some structures might be sterically more exposed to interaction with molecular machinery in the 3D space. Additional structural regions might help in proper folding of these motifs and play an accessory role in maintaining the SINEUP function. Thus, any deletion might inadvertently disrupt functional structures and interaction with RBPs.

We recently reported that deletion of certain regions in antisense *Uchl1* SINEB2 RNA is detrimental to SINEUP function^12^. Several of these deleted regions correspond to the SINEUP structure motifs identified in this study: Δ100-111 disrupts motif 7, Δ130-141 is the deletion of the 3ʹ end side of motifs 3 and 4, and Δ5-14 disrupts motifs 1 and 2^12^. Interestingly, their Δ65–74 mutant^12^, which deletes part of motif 6, does not show a significant decrease in the SINEUP function, whereas another study demonstrated that deletion of SINEB2 RNA SL1 consisting of the entire motif 6 abolishes SINEUP activity^11^. The in-cell structure of human SINE-derived FRAM RNA also showed a partial match for SINEUP motifs and the presence of a structural feature analogous to SL1. Furthermore, we found similarity between the structure of the motif 6 sequence in 2D and that of a motif in 18S rRNA in 3D, and the structure of motif 2 was found to be similar to a region in 16S rRNA in 3D. The matching region in 18S rRNA belongs to helix 34, which forms the roof of the mRNA entry channel. The similarity of the SL1 region of SINEB2 RNA with rRNA was previously noticed in a molecular dynamics simulation based on NMR data^44^, and here in-cell analysis again showed that the conserved and functionally crucial SINEUP motifs were similar to rRNA structure motifs.

Mouse SINEB2s originate from tRNA element^57,61^, and pseudouridylation of tRNA bases is reported to be crucial for its stability and function^35^. Not only tRNAs, but all the RNAs involved in translation contain several Ψ-modified sites^34,35^. This was also found to be the case for SINEUPs in this study. We identified two Ψ modifications at positions 16 and 112 of antisense *Uchl1* SINEB2 RNA. The Ψ at position 112 is inside motif 5, and that at position 16 is just at the motif 2 boundary. This indicates that pseudouridylation of these SINEB2 RNA positions might help to stabilize the structure of these motifs by forming extra hydrogen bonds by N1-H position of Ψ or by base-stacking, which is an intrinsic property of Ψ^34,62^.

We further analyzed the structural dynamics of both SINEB2 and FRAM RNAs in the nucleus and cytoplasm and identified two regions that differ between their nuclear and cytoplasmic structures, with motif 2 and the region between motifs 5 and 6 of SINEB2 RNA slightly more open in the nucleus than in the cytoplasm. Less structured regions were also observed in the nuclear structure of FRAM RNA. We wondered whether these nucleus-specific structures can provide binding sites for proteins involved in nuclear transport. We knew that binding to HNRNPK is crucial for cytoplasmic co-localization of SINEUP-GFP and sense mRNA complex and their association with polysomes^38^. Furthermore, HNRNPK is reported to interact with c-myc IRES and stimulate its cap-independent translation^63^. Since HNRNPK participates in both nuclear transport and translation regulation, we checked if structurally dynamic regions have any effect on its interaction in subcellular compartments. Indeed, we observed distinctive binding patterns of HNRNPK in the nucleus and cytoplasm; in the nucleus, it interacts with short regions in motifs 1, 3, and 4 of SINEB2 RNA, but in the cytoplasm, it interacts with a much longer double-stranded region covering all but one motif. These particular binding patterns could be connected to different functions of HNRNPK in the nucleus and cytoplasm.

In the cytoplasm, the loop region between motifs 5, 6, and 7 of mouse SINEB2 RNA assumes a cloverleaf-like structure, formed from a multiloop where four stems converge. One stem is SL1, which harbors motif 6, another stem consists of motif 7, and a disrupted motif 5 makes the top of the central stem where the Ψ at position 112 is situated at the multiloop branching point. Interestingly, though the nuclear and cytoplasmic structures of FRAM RNA are highly similar, we discerned a similar cloverleaf-like structure. icSHAPE structures are generally enriched for the most stable and dominant structural conformations in the cell, so any alternative short-lived structures might not have been captured. It is fascinating to see that mouse SINEB2 RNA that originated from tRNA and human FRAM RNA that originated from 7SL RNA^61^ share common characteristics: short conserved structural motifs, a cloverleaf-like structure in the cytoplasm, and positive regulation of translation. Previously, we created a minimal functional SINEUP by deleting nucleotide positions 1-30 and 120-167 from SINEB2 RNA, thus keeping only nucleotide positions 31-119 which is the region corresponding to motifs 5, 6, and 7 as well as SL1 and cytoplasmic cloverleaf-like structure^12^. This mutant upregulated GFP mRNA translation and retained about 80% of full length SINEB2 functional activity supporting the significance of this structural region in SINEUP function^12^.

In addition to the intramolecular structure, we were interested to better understand the intermolecular interactions of SINE-derived RNAs with translation machinery. Interaction with 18S rRNA has been reported to be crucial for both linear and circular IRESs^42,43,53^. Knowing that SINEUP-GFP RNA co-sediments with free small subunits of ribosomes and monosomes, we checked direct interaction of SINE RNA with rRNAs by PARIS2 and observed such RNA pairs. The sparsity of SINEUP–rRNA interactions may be due to a limitation of psoralen cross-linking, which has bias for staggered uridines and may not access RNA structures positioned inside complex ribonucleoprotein regions^55^. Alternatively, the efficiency of PARIS2 to capture RNA–RNA interactions may be limited by the level of expression of RNAs as well as by the brief half-lives of such interactions^41,55^. Nevertheless, our PARIS2 data identifies for the first time a direct interaction of SINEUP-GFP (SINEB2) RNA and miniSINEUP-GFP (FRAM) RNA with both 18S and 28S rRNA, which is consistent with previous polysome profiling data^38^. Even though we capture long RNA-RNA interactions, in terms of sequence complementarity we unambiguously identify only short discontinuous stretches of base-pairing between SINE-derived RNA and rRNA. However, this may still be sufficient to influence translation, given that circular IRESs have 7-mer–long complementarity to 18S rRNA^43^ and HCV IRES needs only 3 nucleotide-long Watson-Crick base-pairing with 18S rRNA to induce translation^53^. Moreover, RNA–RNA interactions in 3D depend heavily on non-Watson-Crick base pairing and non-covalent interactions, such as base-stacking and tertiary contacts by purine to minor-groove and tetraloop to tetraloop receptor^64^, which are not easily predicted. Therefore, we decided to take a closer look at the SINE-rRNA interactions.

Notably, we discovered an interaction between a region comprising helix 14 and 15 of 18S rRNA (Int 1) and motif 1 and 2 of SINEB2 RNA. FRAM RNA motifs 1 and 2 along with motifs 5 and 6 also interacted with helix 15 of 18S rRNA. Helix 14 of 18S rRNA is known to interact with ribosomal protein rpL23^65^. In addition, a protein ABCE1 (ATP Binding Cassette Subfamily E Member 1), which is involved in translation initiation and termination, interacts with helix 14 during translation initiation^66^. Moreover, helices 14 and 15 bind with eukaryotic initiation factor 5b (eIF5b), a ribosome-dependent GTPase that mediates joining of 40S and 60S subunits of ribosomes^67^. This indicates the possibility that SINEUP motifs 1 and 2 are involved in the joining of ribosomal subunits during translation initiation. Furthermore, we observed contact of the region between motif 5 and 6 of SINEB2 RNA with domain III helices 55, 56, and 59 of 28S rRNA. Helix 59 is a part of the ribosomal tunnel exit site, from where the nascent peptide emerges out of the ribosome^68^. It suggests that SINEB2 RNA not only takes part in initiating target mRNA translation but also in elongation. In addition, SINEB2 RNA motif 5 and the surrounding region was found to bind with domain I, helix 25, expansion segments ES7c and ES7d of 28S rRNA. The connection with ES7c was also found to be conserved in FRAM RNA, in which the SL1 region (containing a partial match to motif 5) participated in the interaction. ES7 is the largest expansion segment positioned at the back of the 60S ribosomal subunit. This highlights the importance of SINEUP structure motifs in their interaction with the large subunit of the ribosome. We also observed interaction of FRAM RNA SL1 with 18S rRNA helices 21, 37, and 38. Because the helix 38–42 region is crucial for mRNA binding and placement of anticodon arms for tRNAs^69^, this interaction is notable for the proper positioning of SINEUP target mRNA and translation initiation. FRAM RNA also interacted with domains I, III, IV, and V of 28S rRNA.

These contacts at multiple rRNA sites indicate the dynamic changes in SINE RNA–rRNA interaction that occur as translation progresses and highlight the role of SINEUPs in regulating both translation initiation and elongation. Moreover, SINEUP RNAs were found to interact with other protein-coding RNAs and lncRNAs (**Supplementary Table 4**). However, among these RNA partners only *Ank3* (Ankyrin 3) was observed to interact with the SINEB2 RNA region. ANK3 protein links membrane proteins to spectrin-actin cytoskeleton and is involved in neuronal development^70^, but role of *Ank3* mRNA in translation is not known. It is to be noted that although we captured the dynamics of SINE RNA structure and interaction with rRNA and RBP, many of these processes are very transient in nature and it is unlikely to obtain a complete view of all intermediate RNA structures and interaction kinetics in possibly very short time windows by existing in vivo methods. Considering this, further 3D structure analysis of the tertiary connections of SINEUP RNA with rRNA and translation complexes by using cryogenic electron microscopy or other approaches is needed in the future to construct a full 3D model of larger molecular complexes. In this study, the experimental 2D structure data-based prediction of 3D structure models for 18 SINE-derived RNAs identified stabilizing stacking interactions within structure motifs and demonstrated strikingly similar dynamics of mouse and human SINE-derived RNAs in 3D space in the subcellular compartments. We list all the base-stacking and non-canonical interactions determined from these models, which will provide a foundation for future experimental 3D structure analysis. In addition, the 3D structure models of functionally inactive antisense *Gadd45α* SINEB2-a and related structure mutants confirmed the similarity in global structure and SL1 of functional Δ27-36 mutant with antisense *Uchl1* SINEB2 which was missing in the wild-type and other non-functional mutants. This further supports the idea that such short structure motifs are crucial for non-coding RNA function.

In summary, this study offers a repertoire of the sequences, structures, and functional features of several SINE-derived RNA elements for the first time and demonstrates that evolutionarily distant SINE RNAs have a convergent translation upregulatory function driven by common short structure motifs. We show that mouse and human functional SINE-derived RNAs follow a similar pattern of structural dynamics and SINE RNA–RBP interactions are compartment-specific. Our study reveals direct interaction of mouse and human SINE-derived transcripts with human small and large subunit of rRNAs, suggesting a mechanism for SINEUP-mediated post-transcriptional gene regulation. This is the first study to provide a large number of RNA 3D structure models of functional SINE-derived RNAs. Altogether the data presented here is a key step towards filling the void in the structural and functional analysis of SINE-transcribed RNA and will facilitate future studies that aim to resolve the SINEUP mechanism and biomolecular complexes involved.

## Data availability

The icSHAPE sequencing data were deposited in Gene Expression Omnibus (GEO) with accession numbers GSE146407 (for whole cell icSHAPE libraries of SINEUP-GFP) and GSE224534 (for whole cell icSHAPE libraries of miniSINEUP-GFP and nuclear-cytoplasmic fractionated icSHAPE libraries), and GSE243220 (for whole cell icSHAPE libraries of antisense *Gadd45α* SINEB2-a deletion mutants). The PARIS2 sequencing data and SINEUP RNA modification data are accessible through GEO accession numbers GSE224533 and GSE224018 respectively. Nuclear-cytoplasmic fractionated seCLIP sequencing data is available through the GEO accession number GSE227250.

## Acknowledgements

We are grateful to the late Dr. Silvia Zucchelli (Department of Health Sciences, Center for Autoimmune and Allergic Diseases (CAAD) and Interdisciplinary Research Center of Autoimmune Diseases (IRCAD), University of Piemonte Orientale, Novara, Italy) for her valuable suggestions and scientific discussion. This work is dedicated to her memory. We are thankful to Dr. Howard Y. Chang from Standford University for gifting us icSHAPE-probing reagent. We thank Dr. Byron Lee from Chang lab at Stanford University for helpful suggestions for successful icSHAPE library preparation. We thank Souta Nagayama from P. Carninci’s lab for his help in deletion mutant analysis.

## Funding

This work was supported by RIKEN Centre for Integrative Medical Sciences and RIKEN Centre for Life Science Technology from the Japanese Ministry of Education, Culture, Sports, Science and Technology (MEXT); and Basic Science and Platform Technology Program for Innovative Biological Medicine grant from the Japan Agency for Medical Research and Development (AMED grant no.: 18am0301014 to PC).

## Conflict of interest statement

S.G. and P.C. are inventors on patent US9353370B2 and related applications in Europe, USA and Japan, and H.S., H.T., S.G., and P.C. are inventors on patent application 102018000002411 and the related application PCT/IB2019/050914 held by SISSA in Italy and TranSINE Therapeutics in the UK, of which S.G. and P.C. are founders and H.T. owns shares.

H.S., N.T., M.V., H.T., and P.C. are inventors on the E.U. patent application EP3992289A1 (WO2022090733A1) held by TranSINE Therapeutics in the UK. S.G., N.T., H.T., and P.C. are inventors on patent application WO2022064221A1 held by TranSINE Therapeutics in the UK. These conflicts of interest were not associated with the funding of this work.

## Author Contributions

H.S. H.T. and P.C wrote, reviewed, and edited the manuscript with comments from all the authors and interpreted the data. H.S. designed and performed the SINEUP functional analysis, whole cell and nuclear-cytoplasmic fractionated icSHAPE, icSHAPE for deletion mutants and analyzed respective data. H.S. performed 2D and 3D structure modelling and analyzed PARIS data. M.N.Z.V. analyzed seCLIP data and RNA modification data from Oxford Nanopore direct-RNA sequencing. N.T. performed seCLIP, RNA modification experiments and prepared direct-RNA sequencing libraries. H.S.N. prepared PARIS libraries and icSHAPE libraries for deletion mutants. H.T., P.C, and S.G. conceptualized and provided scientific discussion. P.C. provided resources and secured funding for the project.

## Materials and Methods

### Plasmids and cloning

SINEUP-GFP (pcDNA3.1-based) synthetic SINEUP containing antisense *Uchl1* SINEB2^10^ was used to construct long SINEUP effector domain–swapping mutants. To swap the antisense *Uchl1* inverted SINEB2 of SINEUP-GFP with other SINEB2 sequences of interest, two restriction sites for SacII and ClaI restriction enzymes were introduced at the start and end of the SINEB2. For this purpose, a QuikChange II Site-Directed Mutagenesis Kit (Agilent, catalog #200523) was used. All the mutagenesis primers were designed using QuikChange Primer Design Program (Agilent) and are listed in **Supplementary Table 5a**. The SINEB2 regions were PCR-amplified from RIKEN mouse cDNA clones AK143784 (antisense to *Txnip*) and AK054076 (antisense to *Gadd45α*), then digested by the SacII and ClaI restriction enzymes and sub-cloned in SINEUP-GFP to create swapping mutants of antisense *Txnip* and antisense *Gadd45α* SINEB2-a and -b respectively. Antisense *Gadd45α* SINEB2-ab and ba, and all the full constructs of antisense *Uxt* SINEB2–swapping mutants were obtained from Genewiz. The highly active antisense *Uchl1*_Δ5ʹ-32nt mutant^30^ was included in screening as a positive control. Direct and deleted antisense (ΔAS) *Uchl1* SINEB2 mutants were created by a conventional restriction digestion, PCR, and cloning method, and the SINEUP mutant with deleted binding domain (ΔBD) was purchased from TransSINE Technologies.

Previously described miniSINEUP-GFP^29^ plasmid consisting of antisense *Uchl1* SINEB2 as the effector domain and −40/+4 binding domain design with respect to the start codon of GFP mRNA was used as a backbone for miniSINEUP effector-domain swapping. A conventional PCR amplification, restriction digestion, ligation, and cloning strategy was used to generate miniSINEUP-GFP mutants shown in **Fig. 1e**. The SINEB2 region of RIKEN mouse cDNA clones AK029359 (from 774–960 bp for antisense *Uxt-b* SINEB2), AK032194 (from 769–982 bp for antisense *Nars2* SINEB2), AK041742 (from 2334–2533 bp for antisense *Abhd11* SINEB2), and AK048309 (from 1203–1416 bp for antisense Epb4.9 SINEB2) was amplified by PCR (all primer sequences are shown in **Supplementary Table 5b**). To prepare the effector domain for antisense *Uchl1* SINEB2 + antisense *Uxt-b* SINEB2 (**Fig. 1e**), the SINEB2 regions of RIKEN cDNA clones AK078321 (from 521–690 bp for antisense *Uchl1* SINEB2) and AK029359 (from 774–960 bp for antisense *Uxt-b* SINEB2) were first amplified by PCR separately and then ligated to make one effector-domain insert (all primer sequences are shown in **Supplementary Table 5b**). Similarly, in the case of 3× antisense *Uchl1* SINEB2, PCR-amplified SINEB2 from AK078321 (from 521–690 bp) was consecutively ligated twice to generate three repeats of the same SINEB2. The rest of the SINEB2 effector domains originating from various mouse natural antisense lncRNAs (**Fig. 1e**, and **Supplementary Table 1**) and the consensus sequence for SINEB2 subfamily B3 from the Repbase database^27,47^ were commercially obtained from Hokkaido System Science Co., Ltd. The backbone plasmid, miniSINEUP-GFP, and amplified/commercial SINEB2 effector-domain inserts were digested by EcoRI and HindIII restriction enzymes, then ligated and cloned.

The commercially available pEGFP-C2 (Clontech) plasmid was used as a sense EGFP construct. The miniSINEUP-GFP FRAM plasmid design has been previously described^31^.

### Cell culture and transfection

Human embryonic kidney (HEK293T/17) cells (CRL-11268, ATCC) were grown in a 10-cm culture dish in DMEM (1×) + GlutaMAX-1 (Thermo Fisher Scientific, 10569-044) supplemented with 10% fetal bovine serum (Sigma, 172012) and 1% penicillin streptomycin solution (Wako, 168-23191) at 37°C and 5% CO_2_ for 3–5 days. 70–90% confluent cells were treated with 0.05% w/v Trypsin-0.53 mmol/L EDTA 4Na Solution with phenol red (Wako, 202-16931) for 5 min at 37°C to detach the cells, and 0.5 × 10^6^ cells were seeded in each well of 6-well plates (Falcon). After 24 h, cells were transfected with pEGFP-C2 and SINEUP-GFP (long) or miniSINEUP-GFP swapping mutant plasmids at a molar ratio of 1:4.3 for long SINEUPs GFP [0.19 pmol pEGFP-C2 + 0.81 pmol of long SINEUP] and 1:4.6 for miniSINEUP-GFP [0.19 pmol pEGFP-C2 + 0.87 pmol of miniSINEUP-GFP] using 10 µL of Lipofectamine2000 (Thermo Fisher Scientific, 11668-019). After 24 h of transfection, cells were harvested.

### Protein extraction and Western blot

Cells were washed with D-PBS (-) (Wako, 045-29795) and lysed using 140 µL of 10× Cell Lysis buffer (Cell Signaling Technology, 9803S) with PMSF (Cell Signaling Technology, 8553S) per well in a 6-well plate. Samples were rotated at low speed for 1 h at 4°C and then centrifuged (20,000 × *g*) for 10 min at 4°C. The supernatant was collected, and the protein concentrations were measured by DC Protein Assay (BioRad, 500-0116). The absorbance (750 nm) was measured by Multimode Plate Reader ARVO X3 (PerkinElmer). For Western blotting, 20 µg of extracted proteins mixed with 2× SDS sample buffer were separated on 10% SDS PAGE gel (Mini PROTEAN TGX Precast Gel, 10%, 12-well comb; BioRad, 456-1035) and transferred to a nitrocellulose membrane (Amersham Protran Premium ECL 0.45 µm; GE Healthcare Life Sciences, 10600013) through semi-dry transfer at 25 V for 30 min using Trans-BLOT SD Semi-dry Transfer Cell (BioRad). Tris-glycine buffer containing 20% methanol was used for transfer. The membranes were blocked with 5% nonfat dry milk (Cell Signaling Technology, 9999S) in Tris Buffered Saline with Tween-20 (10× TBST, Cell Signaling Technology, 9997) for 30 min with shaking. Proteins were immunoblotted with primary antibodies followed by horseradish peroxidase– (HRP) conjugated secondary antibodies. EGFP was detected by anti-GFP rabbit serum A-6455 (Life Technologies), ACTINB was detected by monoclonal anti-β-actin antibody (Sigma Aldrich, A5441). Protein signals were detected by ECL Western Blotting Detection Reagent (GE Healthcare Life Sciences, RPN2109) with FUJI LAS-3000 system (Fujifilm) and FUSION (Vilber-Lourmat). The band intensities were analyzed by Image J version 1.48 software (National Institutes of Health).

### RNA extraction, cDNA synthesis, and quantitative RT-PCR

HEK293T/17 cells were washed with D-PBS (-) (Wako, 045-29795) and detached using 0.05% w/v trypsin (Wako, 202-16931). Cells were collected by centrifugation for 5 min (6000 × *g*) at 4°C, and total RNAs were subsequently extracted using RNeasy Mini Kit (Qiagen, 74104). Extracted RNA samples were processed through three rounds of DNase I digestion using a TURBO DNA-free Kit (Invitrogen, AM1907). RNA quality was checked on Agilent 2100 Bioanalyzer (Agilent Technologies) using an Agilent RNA 6000 Nano Kit. Then, 500 ng of the total RNA was used for cDNA synthesis using the Prime Script 1st Strand cDNA Synthesis Kit (TaKaRa, 6110A). Random 6-mer primers (0.3 µL) and 1.7 µL of oligodT primers were mixed to capture RNAs. 2× diluted cDNA (1 µL) was used as a template for the quantitative real-time PCR (qRT-PCR). For qRT-PCR, a SYBR Premix Ex Taq (Tli RNaseH Plus) Kit (TaKaRa, RR420S) was used. Each reaction contained 1 µL of cDNA, 0.4 µL of reverse primer (10 µM), 0.4 µL of forward primer (10 µM), 10 µL of SYBR premix ExTaq, 0.4 µL of ROX reference dye (50×), and 10 µL of nuclease-free water (reaction volume was 20 µL). EGFP and SINEUP-GFP levels were normalized by GAPDH. qRT-PCR was performed by StepOnePlus and 7900HT Real-Time PCR Systems (Applied Biosystems) under the following conditions: hold stage (1 cycle) for 30 s at 95°C, cycling stage (40 cycles) for 5 s at 95°C and for 30 s at 60°C, melt curve stage. Reverse transcription samples were taken to check for any plasmid DNA contamination in cDNA. Each sample was performed in technical triplicates (n = 3, Cт standard deviation > 0.2) and with biological replicates (n ≥ 3). 2^-ΔΔCt ±SD was analyzed by StepOne v2.3 and SDS v2.4 software (Thermo Fisher Scientific). The qRT-PCR primer sequences were as follows:

hGapdh_Fw: TCTCTGCTCCTCCTGTTC

hGapdh_Rv: GCCCAATACGACCAAATCC

EGFP_Fw: GCCCGACAACCACTACCTGAG

EGFP_Rv: CGGCGGTCACGAACTCCAG

SINEUP-GFP_Fw: CTGGTGTGTATTATCTCTTATG

SINEUP-GFP_Rv: CTCCCGAGTCTCTGTAGC

miniSINEUP-GFP_Fw: TTAATACGACTCACTATAGGGAGAC

miniSINEUP-GFP_Rev: CCGCATCTAGTGAACCGTCA

### Phylogenetic analysis and hierarchical clustering

MEGA version 5 software^71^ was used for sequence-based alignment, and mouse SINEB2 sequences tested as effector domains in long and mini SINEUPs were aligned by ClustalW. SINEB2 sequence–based molecular phylogenetic analysis was done using the Maximum Likelihood method. The evolutionary history was inferred by the Maximum Likelihood method based on the Tamura 3-parameter model. Initial tree(s) for the heuristic search were obtained automatically by applying Neighbor-Join and BioNJ algorithms to a matrix of pairwise distances estimated using the Maximum Composite Likelihood approach and then selecting the topology with superior log likelihood value. A discrete Gamma distribution was used to model evolutionary rate differences among sites (5 categories (+G, parameter = 4.1774)). The tree was drawn to scale, with branch lengths measured in the number of substitutions per site.

### icSHAPE library preparation

The icSHAPE standard protocol was followed^12,49^. Briefly, HEK293T/17 cells were co-transfected with pEGFP-C2 and SINEUP-GFP or miniSINEUP-GFP swapping mutants using Lipofectamine2000 reagent in 6-well plates. Cells were harvested after 24 h of transfection, pooled together, and treated with NAI-N_3_ reagent (modified) (NAI-N_3_ reagent was received as a gift from Dr. Howard Chang’s lab at Stanford University) for in vivo SHAPE modification or with DMSO (mock control). Two biological replicates for each of the NAI-N_3_ and DMSO libraries were made. RNA was extracted using RNeasy Mini Kit (Qiagen, 74104) and digested with DNaseI to remove DNA contamination using a TURBO DNA-free Kit (Invitrogen, AM1907). RNA was ribo-depleted using a magnetic RiboZero kit (Illumina, MRZH11124), and RNA quality was checked by Agilent 2100 Bioanalyzer (Agilent Technologies) nano and pico chip. Next, 500 ng of ribo-depleted RNA was biotin-clicked to label-probed bases followed by RNA fragmentation for 40 s using RNA fragmentation reagents (Thermo Fisher Scientific, AM8740). Fragmented RNA was end-repaired, and the icSHAPE adapter ligation standard protocol was followed^49^ (adapter sequences in **Supplementary Table 6a**). Next, size selection was performed, and RNA was reverse-transcribed using icSHAPE-specific barcoded reverse-transcription primer followed by streptavidin-based selection and size selection of truncated cDNA products. Purified cDNA was circularized using CircLigase II enzyme (Epicentre, CL9025K), then library PCR, size selection, and quantification were performed. Resultant libraries were sequenced on a HiSeq2500 platform (Illumina, 100 bases, single-end). All the oligos used in the icSHAPE library preparation are noted in **Supplementary Table 6a**.

### Comparison of icSHAPE profiles, secondary structure modelling, and motif analysis

The standard icSHAPE bioinformatics pipeline was used^49^. Briefly, after removing PCR duplicates and adapter trimming, raw sequencing data were mapped to EGFP and SINEUP transcripts index using Bowtie2. Next, transcript abundance and reverse transcription (RT) stop numbers were calculated before calculating the correlation between replicates and replicate merging. The mean of the RT stops of the 90– 95% most reactive bases was normalized as 1 and all other RT stops were scaled proportionally. Then enrichment reactivity scores were calculated by subtracting the background signal (DMSO data), and outliers were removed by 90% Winsorization (in which the top 5th percentile is set to 1 and the bottom 5th percentile is set to 0). Valid enrichment reactivity scores were filtered and visualized in integrated genome viewer (IGV v2.11.1). The icSHAPE enrichment scores were used as a soft constraint in RNAfold (ViennaRNA Package)^72^. Linear mapping method was used to derive pairing probabilities, and a method described by Zarringhalam *et al.* (2012)^73^ was used to incorporate guiding pseudo energies into the folding algorithm. To draw the secondary structure, RNAcanvas^74^ (https://rna2drawer.app/) was used.

Sequence and icSHAPE-guided 2D structure of all the SINEB2s were pairwise compared with the structure and sequence of antisense *Uchl1* SINEB2 using ExpaRNA tool^50^, which computes the exact matching sequence-structure motifs common to two RNAs. Thus, derived motif files from all the pairwise comparisons were then screened to extract the positions of Uchl1 SINEB2s that were present in one or multiple motifs in all of the comparisons. Since SINEB2 sequences are highly variable, in most cases only a partial sequence match could be found. Due to their repeating sequence composition, in some instances one query sequence was matched to multiple motifs, or one motif matched to multiple positions in the query. In that case, the longer matching stretch was preferred, and different motif numbers were assigned based on their sequence and structure neighborhood and position on SINEB2. Additionally, SINEUP 2D motifs were compared with existing 3D motifs in the RNA Characterization of Secondary Structure Motifs database (CossMos)^51^.

### Nuclear-cytoplasmic fractionation for icSHAPE

HEK293T/17 cells were co-transfected with 0.8 pmol SINEUP-GFP or miniSINEUP-GFP FRAM (0.8 pmol) and 0.2 pmol sense pEGFP-C2 plasmids (Clontech) in each well of a 6-well culture plate. Four 6-well plates for each sample were prepared. At 24 h post-transfection, the medium was aspirated, and the cells were washed and collected in 1 mL of 1× D-PBS (-) (Nacalai tesque, 14249-24) per well. Cells from one 6-well plate (∼1 × 10^7^ cells) were pooled together in a 15-mL tube and centrifuged at 300 × *g* for 2 min at room temperature. The resultant supernatant was removed, and the pelleted cells were resuspended in either 250 µL of 100 mM NAI-N3 reagent (Enamine, EN300-247313) for modification or in 5% DMSO (Sigma Aldrich, D2650) solution in 1× PBS (Gibco) for mock treatment. The reaction was incubated on a rotating wheel at 37°C for 5 min at room temperature and then stopped by centrifugation at 2500 × *g* for 1 min at 4°C and further removal of the supernatant. Next, the protocol by Pandya-Jones and Black (2009)^75^ was utilized for cellular fractionation. In brief, the NAI-N3- and DMSO-treated cell pellets were resuspended in ice-cold NP-40 lysis buffer (10 mM Tris-HCl pH 7.5, 0.15% NP-40, 150 mM NaCl), then the lysate was added on top of 2.5 volumes of chilled 25% sucrose cushion in lysis buffer without NP-40 followed by centrifugation at 19,000 × *g* for 10 min at 4°C. The resultant supernatant containing the cytoplasmic fraction was collected. The pellet was further gently rinsed with 200 µL chilled 1× PBS/1 mM EDTA and centrifuged at 9,730 × *g* for 2 min at 4°C to remove the supernatant. Next, to extract the nuclear fraction, the protocol by Brugiolo *et al.* (2017)^76^ was followed. The pellet was resuspended in 100 µL nuclear buffer I (50% glycerol v/v, 20 mM Tris-HCl pH 7.9, 75 mM NaCl, 0.5 mM NaCl, 0.85 mM DTT, 0.125 mM PMSF) and then in 900 µL nuclear buffer IIA (20 mM HEPES pH 7.6, 300 mM NaCl, 0.2 mM EDTA, 1 mM DTT, 7.5 mM MgCl_2_, 1M Urea, 1% NP-40, 400 U of RNase inhibitor). The solution was mixed thoroughly by vortexing and then pipetting up and down, and then incubated on ice for 20 min. Finally, the nucleoplasmic fraction was extracted by centrifuging at 14,000 × *g* for 5 min at 4°C and collecting the supernatant. The success of fractionation was checked by Western blot and qRT-PCR for known nuclear and cytoplasmic marker genes. Four nucleoplasmic or cytoplasmic fractions, each coming from four 6-well plates of the same sample, were pooled together for further RNA extraction and icSHAPE library preparation as described above. All the steps are the same as in the original protocol^49^, except the removal of free 3’ RNA linkers after ligation, for which the completed ligation reaction was treated with 4 µL RecJ_f_ (New England Biolabs, M0264L) and 3 µL 5ʹ Deadenylase (New England Biolabs, M0331S) at 37°C for 1 h and further cleaned up by using a Zymo RNA Clean and Concentrator-5 purification kit (Zymo Research, R1016). All the oligos used in the library preparation are noted in **Supplementary Table 6a**.

### seCLIP library preparation from nuclear-cytoplasmic fractionated samples

Approximately 0.5 × 10^7^ HEK293T/17 cells were plated on a 10-cm culture dish and after 24 h, transfected with SINEUP-GFP and pEGFP-C2 plasmids as described above. After 24 h of transfection, 2 × 10^7^ cells were gently washed with 1× PBS and then crosslinked by 400 mJ/cm, 254 nm UV light irradiation on ice with the lid off for 15 min. Next, the cells were collected by scraping in 1 mL chilled PBS followed by centrifugation at 200 × *g* for 5 min at 4°C. The resultant supernatant was discarded, and the pellet was used for nuclear-cytoplasmic fractionation following the protocol by Brugiolo *et al.* (2017)^76^. The pellet was first resuspended in 2 mL hypotonic buffer (10 mM Tris-HCl 7.5 pH, 10 mM KCl, 1.5 mM MgCl_2_, 0.5 mM DTT, 1:200 proteinase inhibitor just prior to use) and incubated on ice for 5 min followed by centrifugation at 500 × *g* for 10 min at 4°C. The supernatant containing hypotonic fraction was discarded, and the pellet was resuspended in 1 mL lysis buffer 0.3 (50 mM Tris-HCl 7.5 pH, 150 mM NaCl, 2 mM MgCl_2_, 0.3% NP-40, 1:200 proteinase inhibitor just prior to use) and incubated on ice for 10 min. The mix was then centrifuged at 1000 × *g* for 5 min at 4°C, and the supernatant containing the cytoplasmic fraction was collected in a new 1.5-mL tube. The pellet was mixed with 1 mL lysis buffer 0.5 (50 mM Tris-HCl 7.5 pH, 150 mM NaCl, 2 mM MgCl_2_, 0.5% NP-40, 1:200 proteinase inhibitor just prior to use) and incubated on ice for 10 min followed by centrifugation at 1000 × *g* for 5 min at 4°C. The supernatant was discarded, and the pellet was first resuspended in 100 µL nuclear buffer I (as used for icSHAPE fractionation, without addition of PMSF) and then in 900 µL nuclear buffer IIA (as used for icSHAPE fractionation) with 1:200 proteinase inhibitor just prior to use. The pellet was mixed well by vortexing and then pipetting up and down, and then incubated on ice for 15 min. The reaction mix was then centrifuged at 15,000 × *g* for 5 min at 4°C, and the nucleoplasm containing supernatant was transferred to a new 1.5-mL tube. All fractions were separately centrifuged at 20,000 × *g* for 5 min at 4°C and tested for the presence of markers specific to nuclear or cytoplasmic fractions by Western blot.

Next, the seCLIP protocol described by Van Nostrand *et al*. (2017)^46^ was utilized to prepare nuclear and cytoplasmic seCLIP libraries. In brief, nucleoplasmic and cytoplasmic lysates were treated with 4 U of DNase I and 10 µL of 1:25 diluted RNase I (100 U/µL) in a thermomixer at 37°C and 1200 rpm for 5 min. The reaction was then cooled on ice, 7 μL Murine RNase Inhibitor (New England Biolabs, M0314) was added, and the mix was centrifuged at 15,000 × *g* for 15 min at 4°C. The resultant supernatant was mixed with 15 µg of anti-hnRNPK antibody (Abcam, ab39975) coupled to 200 µL magnetic beads (Dynabeads M-280 Sheep Anti-mouse IgG, Invitrogen, 11201D) and incubated at 4°C overnight. Around 2% of the input samples (containing all RNA-protein complexes) were saved before immunoprecipitation. Immunoprecipitated samples were first end-treated with FastAP thermo-sensitive alkaline phosphatase (Thermo Scientific, EF0651) and T4 poly nucleotide kinase (New England Biolabs, M0201L) and then ligated with 3ʹ RNA linker (with sample barcodes, see sequence in **Supplementary Table 6b**). Next, both immunoprecipitated and input RBP–RNA complexes were visualized by Western blot imaging and the region corresponding to HNRNPK and above (from 55–180 kDa) was sliced from the preparative nitrocellulose membrane blot. The RNA was extracted by treating the membrane slices with urea and proteinase K followed by acid phenol–chloroform extraction and purification by using a Zymo RNA Kit (R1013). Further treatment of samples and cDNA library preparation are the same as described in the original protocol^46^. Libraries were sequenced on the HiSeq2500 platform (150 cycles, single-end).

### Analysis of nuclear-cytoplasmic fractionated seCLIP data

Fastq files were processed with two rounds of cutadapt (which trims off adapters and controls for double ligation events) before being mapped to a custom genome (hg38 + SINEUP construct) using STAR v2.7.8a with parameters as in the ENCODE eCLIP project pipeline (https://www.encodeproject.org/eclip/) (namely, --outFilterMultimapNmax 1, --outFilterMultimapScoreRange 1, --outFilterScoreMin 10, --alignEndsType EndToEnd). Reads mapped on the sense strand of the SINEUP construct were removed. Peaks were called on mapped reads using clipper v0.2.0 (https://github.com/YeoLab/clipper) with custom data files (gff, exons bed file) containing the SINEUP region, with default parameters. Scripts from the merge_peaks github repository (https://github.com/YeoLab/merge_peaks) were used to normalize seCLIP reads over input reads for each replicate (overlap_peakfi_with_bam.pl) before merging input-normalized peaks into a final set of significant binding regions (compress_l2foldenrpeakfi_for_replicate_overlapping_bedformat.pl).

### Preparation of chemically modified SINEUP-GFP RNA transcribed in vitro

Transcribed SINEUP-GFP RNA was synthesized in vitro as described by Toki *et al.* (2020)^77^ by using a mMESSAGE mMACHINE SP6 Transcription Kit (Thermo Fisher Scientific) and was then modified by following the protocol by Mandal and Rossi (2013)^78^. For the 20% Ψ modification, UTP was replaced with 20% pseudouridine-5ʹ-triphosphate (TriLink, USA; final concentration, 7.5 mM) by mixing the reagents with 40 ng/µL of linearized SINEUP RNA. To add a poly-A tail, 1–10 µg of RNA transcribed in vitro was treated with Escherichia coli poly(A) polymerase (5000 U/mL, New England Biolabs, M0276) at 37°C for 30 min followed by clean-up using an RNeasy Mini kit (Qiagen) and the modified transcripts of SINEUP-GFP RNA were recovered.

### Extraction of modified SINEUP-GFP RNA transcribed in-cell

HEK293T/17 cells were plated on a 10-cm dish and SINEUP-GFP plasmids were transfected for 24 h; then 2 × 10^7^ cells were harvested by Trizol/chloroform extraction, and total RNAs containing SINEUP-GFP RNAs at the aqueous phase were extracted by using an RNeasy mini Kit (Qiagen). The RNA was then digested with 4 U of TURBO DNase (TURBO DNA-free kit, Invitrogen, AM1907) at 37°C for 90 min followed by DNase inactivation.

### Pull-down of SINEUP-GFP RNAs transcribed in-cell

The total RNA volume was adjusted to 100 µL with lysis buffer (50 mM Tris-HCl pH 7.0, 10 mM EDTA, 1% SDS, 1:100 RNase inhibitor) and mixed with two volumes of hybridization buffer (750 mM NaCl, 50 mM Tris-HCl pH 7.0, 1 mM EDTA, 1% SDS, 15% formamide, 1:200 RNase inhibitor) with additional 6.6% formamide. Then, 100 pmol of SINEUP-GFP-specific total probe (2 µL of 50 µM probe) was added to this mix (probe design as described by Toki *et al.*, 2020)^38^, and the reaction was incubated at 37°C overnight with vertical rotation. Next, 100 µL of MagCapture Tamavidin 2-REV magnetic beads (WAKO) were washed and resuspended in 100 µL lysis buffer. This bead suspension was added to the completed hybridization reaction and incubated at 37°C for 30 min with mixing. The solution was then magnetically separated, and the supernatant was discarded. The beads were washed four times with pre-warmed wash buffer (2× saline sodium citrate, 0.5% SDS, 1:200 RNase inhibitor), and the supernatant was discarded. After this, the beads were resuspended in 100 µL ProK buffer (10 mM Tris-HCl pH 7.5, 0.8% SDS, 15 mM EDTA) without proteinase K and incubated at 65°C for 5 min with end-to-end mixing at 450 rpm. The first pull-down RNA was then extracted by the Trizol/chloroform method and purified by miRNeasy mini column (Qiagen, 217004). This pull-down and RNA extraction was repeated once again to enrich SINEUP-GFP RNAs.

### Library preparation of Oxford Nanopore direct-RNA sequencing

Libraries of the direct SINEUP-GFP RNA transcribed in vitro and in-cell were prepared with an SQK-RNA002 kit (Oxford Nanopore Technology). The original reverse-transcription adapter (RTA) was used for in vitro samples and four barcoded RTA adapters described by Leger *et al.* (2021)^79^ were used for in-cell samples. The libraries were applied to MinION Mk1C sequencer (Oxford Nanopore Technologies) and sequenced for 72 h.

### Modification detection using long-read sequencing

Raw fast5 files from direct-RNA sequencing of modified SINEUP RNA transcribed in-cell and in vitro were base-called using guppy v5.0.7 with RNA-centric options (--trim_strategy rna, --u_substitution true, --reverse_sequence true). After preparing reference transcriptome annotation files (bed and fasta annotation files from bedparse gtf2bed and bedtools getfasta, respectively), the base-called data were mapped to the reference using minimap2 v2.17. To detect modified k-mers in the data we followed the nanocompore pipeline (https://github.com/tleonardi/nanocompore/). Briefly, this involved indexing the raw data, followed by running nanopolish eventalign to realign the raw signal-level data to the k-mers of the reference. After this, the signal was collapsed to the k-mer level using NanopolishComp, and nanocompore was used to compare the sample transcribed in-cell with the different transcription conditions in vitro. Results of this comparison were filtered by both log odds ratio (absolute value > 1) and *p* value (<0.01). Finally, a peak-calling algorithm (https://gist.github.com/tleonardi/0bb31e6a380e5766f04f4e197d36b38e) was used to get rid of multiple adjacent significant sites, because a single modified nucleotide can often cause multiple neighbouring k-mers to be measured as significant sites, when in reality there is just one modification. The peak-calling algorithm merges these adjacent sites into a single k-mer, which helps to highlight true-positive sites.

### PARIS library preparation

HEK293T/17 cells were transfected with pEGFP-C2 and SINEUP-GFP (SINEB2) or miniSINEUP-GFP FRAM plasmids in 6-well plates as described above. After 24 h of transfection, cells were gently washed with 1× PBS and treated with 150 µL 1× PBS (control) or 1 mg/mL Amotosalen (MedChemExpress, HY-107004A) crosslinking solution added to each well of the 6-well plates, followed by incubation under standard cell-culture conditions (37°C, 5% CO_2_) for 30 min. Next, cells were irradiated for 30 min by 365-nm UV light while on ice with swirling of the plate every 10 min. Then, the crosslinking solution was removed, and the cells were scraped off into 1 mL chilled 1× PBS in each well and transferred to a 15-mL tube. Crosslinked cells were pelleted by centrifuging at 300 × *g* for 5 min at 4°C, and the supernatant was discarded. Next, for the total nucleic acid (TNA) extraction and PARIS library preparation, the protocol by Zhang *et al.* (2021)^41^ was used with some modifications. Briefly, the cell pellet was lysed by 6 M GuSCN with pipetting followed by treatment with 0.5 M EDTA and passed through a 25G needle 20 times. After this, the proteins were digested by proteinase K (final concentration 1 mg/mL) at 37°C for 1 h on a thermomixer with shaking, and solution was again passed through the 25G needle. After TNA was precipitated by phenol-isopropanol precipitation as described in Zhang *et al.* (2021)^41^, 100 µg of TNA sample was treated with 50 units of TURBO DNase (Invitrogen, AM2239) at 37°C for 20 min. RNA was precipitated using phenol-isopropanol and further purified using RNA clean and concentrator kit (Zymo Research, R1015). The RNA quality was measured with Bioanalyzer Nano Kit (Agilent). DNase-treated RNA (10 µg) was fragmented by using 5 units of ShortCut RNase III (New England Biolabs, M0245) for 5 min at 37°C and the reaction was stopped by adding 10 µL nuclease free water, 100 µL of RNA binding buffer (Zymo Research, R1015) and 150 µL of 100% ethanol and then purified through a column from the purification kit (Zymo Research, R1015) followed by quality check with a Bioanalyzer small RNA kit (Agilent).

For 2D gel purification of RNA duplexes, 8% 1-mm thick denatured first-dimension gel was prepared using the UreaGel System (National Diagnostics, EC-833). A 10-µL sample of fragmented RNA was run on the first-dimension gel in 0.5x Novex TBE running buffer (Invitrogen, LC6675) at 30 W for 9 min. The gel was visualized under 300 nm transillumination after staining with SYBR Gold (Invitrogen, S11494) and each lane was sliced between 50 bp to topside from the first-dimension gel. Next, 16% 1-mm thick denatured second-dimension gel was prepared using the UreaGel System (National Diagnostics, EC-833). The second-dimension gel was poured to encapsulate the gel slices from the first-dimension gel, and 0.5x TBE buffer warmed to 60°C was used to facilitate the denaturing of crosslinked RNA. The gel was run at 30 W for 50 min and then stained with SYBR Gold, and crosslinked RNA was sliced out from the upper diagonal of the 2D gel. The gel slices were cut into smaller pieces and transferred to a 0.5-mL tube with a hole drilled in the bottom with a 20G needle. This 0.5-mL tube was placed inside a 2-mL tube and centrifuged at 16,000 x *g* for 5 min to crush the gel through the hole. Then 1.5 mL of crushing buffer (20 mM Tris-HCl pH 7.5, 0.25 M sodium acetate, 1 mM EDTA, 0.25% SDS) was added to the crushed gel and rotated at room temperature overnight. After this, 500 µL of gel slurry was transferred to a Micro Spin column (Cytiva, 27356501) and centrifuged at 3400 x *g* for 1 min or until all gel slurry was filtered. This filtered gel was then passed through an Amicon Ultra 10K column (Millipore, UFC501096) and further purified using the Zymo RNA Clean and Concentrator-5 column.

Next, in order to perform proximity ligation, 10 µL of the purified double-stranded RNA was treated with 7.5 unit/µL of T4 RNA Ligase I, high concentration (New England Biolabs, M0437) in a reaction mix containing 1× T4 RNA Ligase Buffer (New England Biolabs, M0437M), 1U SUPERase•In RNase inhibitor (Invitrogen, AM2694), and 0.5 mM ATP at room temperature overnight. The proximity-ligated RNA was purified by using a Zymo RNA clean and concentrator-5 column.

To reverse crosslink the RNA, the samples were mixed with 3 µL of 25 mM acridine orange on ice and irradiated with 254-nm UV light for 30 min. The reverse crosslinked sample was transferred to a new 1.5-mL tube and purified using a Zymo RNA Clean and Concentrator-5 column. Further, the reverse crosslinked RNA was heated at 80°C for 90 s and then snap-cooled on ice. The adapter was ligated by adding 14 μL of adapter ligation mix (1.5 units/µL T4 RNA ligase I high concentration, 1.5 µM adapter (5ʹ Biotin), 10% DMSO, 12.5% PEG 8000, 5 mM DTT, 1× T4 RNA Ligase buffer) to 6 µL of reverse crosslinked RNA and incubating the reaction for 3 h at room temperature. The sequences of all the oligos used in the PARIS library preparation are listed in **Supplementary Table 6c**. After adapter ligation, free adapters were removed by adding 3 units of RecJ_f_ (New England Biolabs, M0264) and 1.7 units of 5ʹ deadenylase (NEB, M0331S) and incubating at 37°C for 1 h. The adapter-ligated RNA was purified using a Zymo RNA Clean and Concentrator-5 column and eluted in 10 µL of nuclease free water. Next, 2 µL of custom reverse transcription primer (with barcode, 1 µM, sequence in **Supplementary Table 6c**) and 1 µL of 10 mM dNTPs were added to the purified RNA and the sample was heated at 65°C for 5 min in a thermal cycler followed by rapid cooling on ice. To this mix, 8 µL reverse transcriptase mix was added, which contained 4 µL of 5x SSIV buffer (Mn2+) (250 mM Tris-HCl, 375 mM CH3COOK, 7.5mM MnCl_2_), 10 mM DTT, 1 unit/μL SUPERaseIn RNase inhibitor (Invitrogen, AM2694), 10 units/µL SuperScript IV (Invitrogen, 18090010). The reaction was incubated at 25°C for 15 min, 42°C for 10 h; hold at 10°C.

The reverse transcribed RNA was captured on streptavidin-coated magnetic beads. For this purpose, 10 µL of Dynabeads M-270 Streptavidin (Invitrogen, 65305) were washed in 20 µL LiCl buffer (7 M LiCl, 20 mM Tris-HCl (pH7.5), 2 mM EDTA, 0.1% Tween20) and then resuspended in 45 µL of LiCl buffer. This beads suspension was added to the reverse transcribed RNA and incubated at 37°C for 30 min with pipette mixing in between. The supernatant was removed by separation on a magnetic stand, and beads were washed three times with 100 µL TE wash buffer (10 mM Tris-HCl (pH7.5), 1 mM EDTA, 0.1% Tween20) and finally resuspended in 30 µL elution buffer (1x RNaseONE buffer, 1% Tween20, 10 µM P3-tall primer). Next, the beads solution was heated at 95°C for 5 min then immediately cooled on ice and magnetically separated to recover the released cDNA containing supernatant in a new 1.5-mL tube. The beads were again resuspended in 30 µL elution buffer, separated on a magnet, and supernatant was recovered and transferred to the released cDNA tube from the previous step. Next, the cDNA was treated with 1 µL of RNaseONE (Promega, M4261), 1 µL of RNaseH (TaKaRa, 2150A), and 1 µL of RNase cocktail (Invitrogen, AM2286) for 30 min at 37°C and further purified by using a Zymo DNA Clean and Concentrator column (Zymo Research, D4033).

After this, 4 µL of circularization reaction mix containing 5 units/µL CircLigase II (Lucigen, CL9021K), 1× CircLigase II buffer (Lucigen, CL9021K), and 2.5 mM MnCl_2_ was added to 16 µL of cDNA and incubated at 60°C for 100 min followed by 80°C for 10 min. Circularized cDNA (1 µL) was used to determine the PCR cycles required for the library amplification. For this, 39 µL of qPCR mix containing 1× Phusion mix (New England Biolabs, M0531S), 0.125 µM P6 Tall v4 primer, 0.125 µM P3 Tall v4 primer, 0.15 µL ROX, 1/3800 SYBR (Invitrogen, S-7563) was added to 1 µL circularized cDNA and amplified at 95°C for 45 s; 20 cycles of 95°C for 15 s, 65°C for 30 s, 72°C for 45 s on a StepOnePlus Real-Time PCR System (Applied Biosystems, 4376598). The qPCR was stopped just before the amplification plot reached the plateau and the corresponding cycle number was noted. For the final library amplification, 6 µL of circularized cDNA was added to 44 µL PCR mix (1× Phusion mix, 0.15 µM P6 Tall v4 primer, 0.15 µM P3 Tall v4 primer) and amplified at 95°C for 45 s; pre-determined cycles of 95°C for 15 s, 65°C for 30 s, 72°C for 45 s on a routine thermocycler. The PCR product was purified using Zymo DNA clean and concentrator column (Zymo Research, D4013) and ran on 6% native TBE gel. DNA from 175 to 300 bp was excised, crushed, and purified. Library quality was checked by using a Bioanalyzer High-sensitivity DNA chip and concentration was quantified by using a qPCR KAPA library quantification kit (KAPABiosystems, KK4835). Four barcoded libraries were pooled together and sequenced on HiSeq2500 platform (150 nt, single-end, rapid mode).

### PARIS data analysis

To check the SINEUP regions interacting with rRNAs, the PARIS2 analysis strategy was used as described by Zhang *et al.* (2021)^41^ (https://github.com/minjiezhang-usc/PARIS2). In short, the 3ʹ end adapters were removed by Trimmomatic v0.39, and the sequencing reads were split according to the sample barcodes using splitFastq script from icSHAPE pipeline^49^. Next, PCR duplicates were removed using the readCollapse script from the icSHAPE pipeline^49^ followed by removal of the 5ʹ header using Trimmomatic v0.39. The reads were then mapped to a custom genome generated by adding SINEUP-GFP (SINEB2), miniSINEUP-GFP (FRAM), and sense EGFP genes as separate chromosomes to the hg38 masked genome^41^, which were previously engineered to contain a single copy of multicopy genes such as rRNA and snRNAs. Mapping was done by STAR v2.7.10a following the parameters described in the PARIS2 paper^41^. To extract SINEUP-rRNA interactions, first the reads that were mapped to rRNA were extracted by samtools (as in the PARIS2 analysis strategy) and then rRNA interacting reads in which one of the arms of the RNA-rRNA pair was mapped to SINEUPs were extracted by using the grep command. The extracted file was sorted by samtools and visualized on the integrated genome viewer (IGV v2.11.1). VfoldMCPX^80^ was used to predict the structure of the RNA-rRNA pair. Human 18S rRNA structure (ID: URS00005A14E2) and 28S rRNA structure (ID: URS0000C873C2) were taken from the RNAcentral database (https://rnacentral.org/).

### 3D structure modelling

SINE RNAs 2D structure information derived from icSHAPE was machine translated into 3D structure models by RNAComposer^81^ (https://rnacomposer.cs.put.poznan.pl/). The 3D models were visualized by using Mol* tool^82^ of RCSB PDB^83^ (https://www.rcsb.org/) and analyzed by RNApdbee 2.0^84^ (http://rnapdbee.cs.put.poznan.pl/) to recognize the positions with base-stacking and non-canonical interactions.

---

**Extended Data Figure 1.**
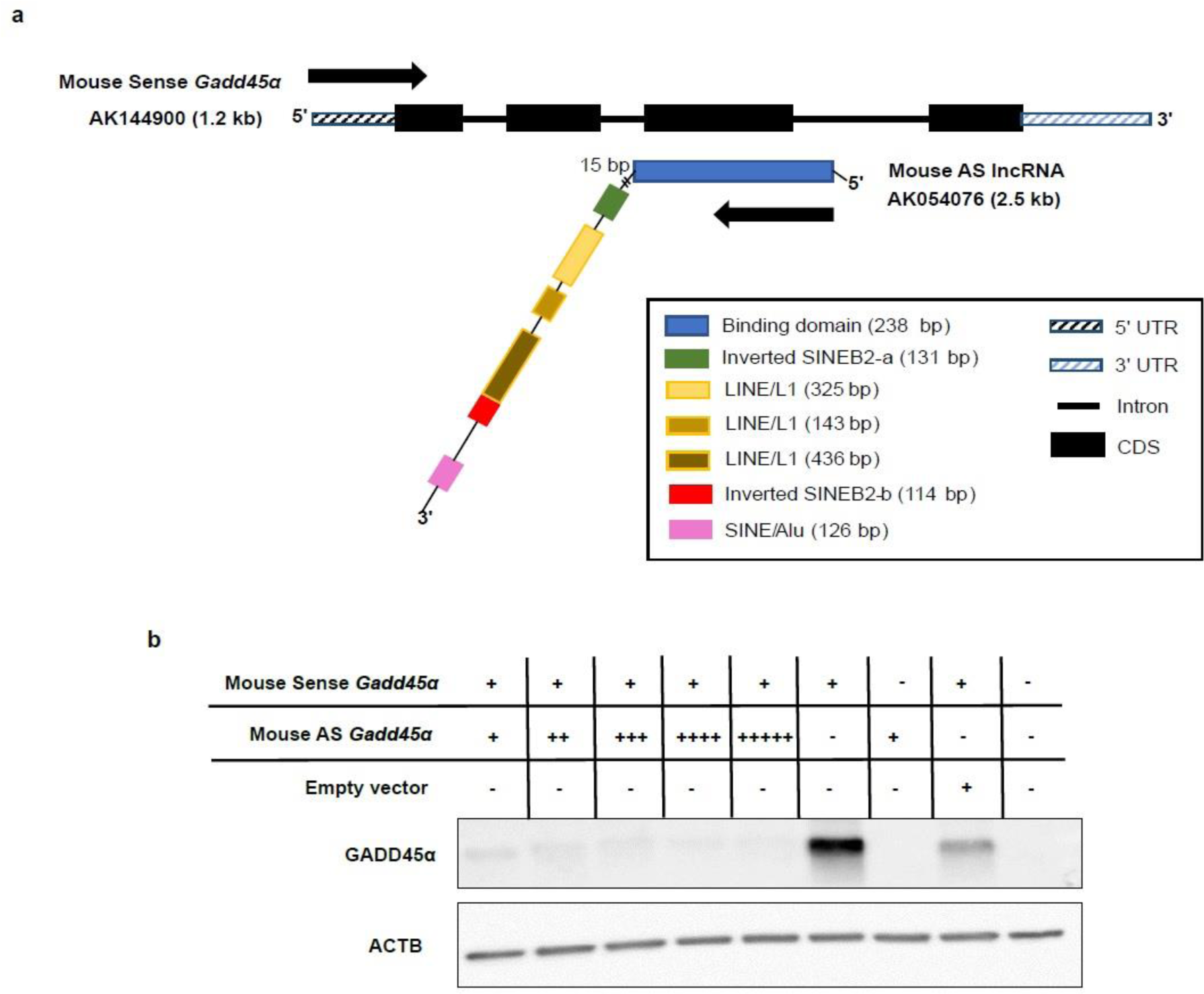
Effect of mouse natural antisense *Gadd45α* lncRNA on sense *Gadd45α* protein expression. (**a**) Molecular features of mouse sense/antisense *Gadd45α* RNAs. Antisense (AS) *Gadd45α* contains a sequence complementary to intron 2, exon 3, and intron 3 of sense Gadd45α mRNA,two inverted SINEB2s, three LINE, and one SINE-Alu elements. (**b**) Western blot analysis of sense Gadd45α protein expression in HEK293T/17 cells co-transfected with sense/antisense *Gadd45α* for 24 h. Empty vector, negative control; plasmid transfection at + 0.7 pmol; ++ 0.8 pmol; +++ 0.9 pmol; ++++ 1.0 pmol; - not transfected. ACTB is the loading control.

**Extended Data Figure 2.**
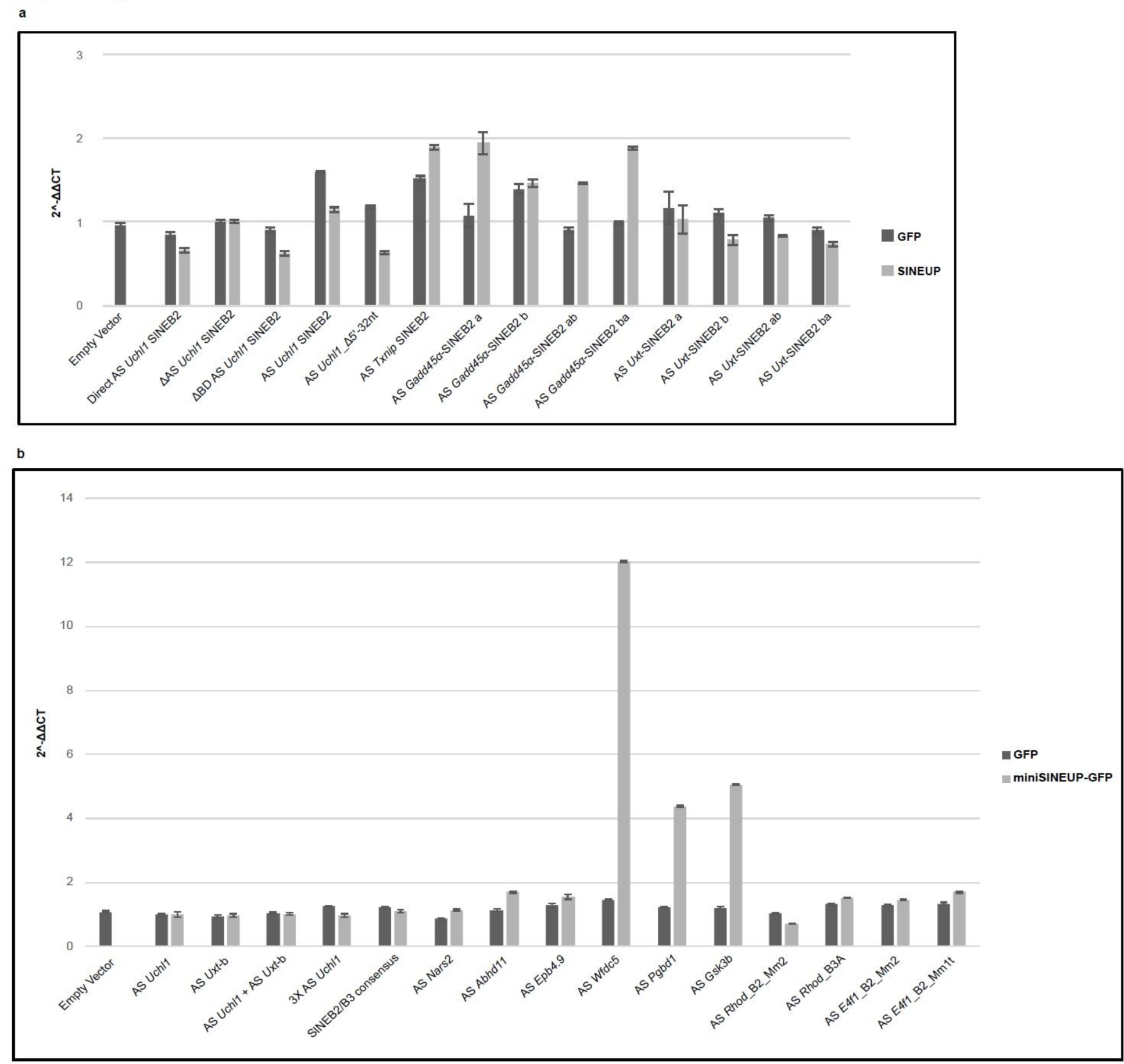
qRT-PCR results for GFP mRNA (black bars) and SINEUP-GFP RNA (grey bars) expression. Expression values are normalized to human GAPDH mRNA. Data were analyzed by using the ΔΔC_T_ method. RNA was extracted from (**a**) long SINEUPs (see description of terms in Figure 1c), with ΔAS *Uchl1* SINEB2 = 1 and from (**b**) MiniSINEUP-GFP (described in Figure 2b), with AS *Uchl1* = 1. Error bars are ± standard error mean for three biological replicates. AS, antisense.

**Extended Data Figure 3.**
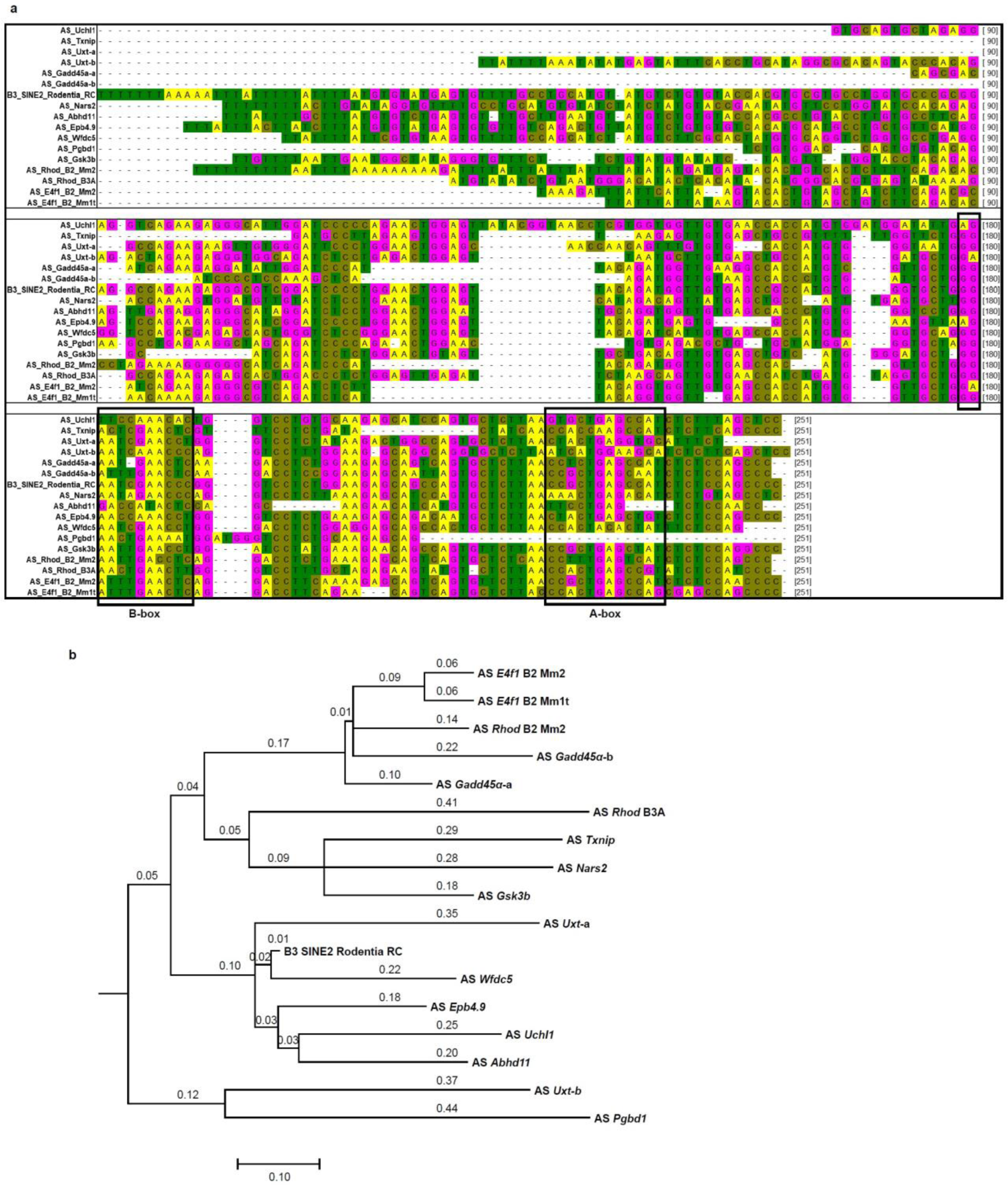
Multiple sequence alignment and phylogenetic analysis of SINEB2 RNA sequences. (**a**) ClustalW sequence alignment of mouse SINEB2 RNA sequences used as effector domain in long and mini SINEUPs. Gaps in the alignment are indicated by -. Bases are highlighted in four different colors. Black squares denote A and B boxes. Alignment is showing the sequence variation within members of the same family of SINE. (**b**) SINEB2 sequence alignment–based molecular phylogenetic analysis by the Maximum Likelihood method (Tamura 3-parameter model) to infer the evolutionary history. The tree with the highest log likelihood (−2415.7806) is shown. The tree is drawn to scale, with branch lengths measured in the number of substitutions per site. AS, antisense; B3 SINE2 Rodentia RC, the consensus sequence for SINEB2 subfamily B3 from the Repbase database.

**Extended Data Figure 4.**
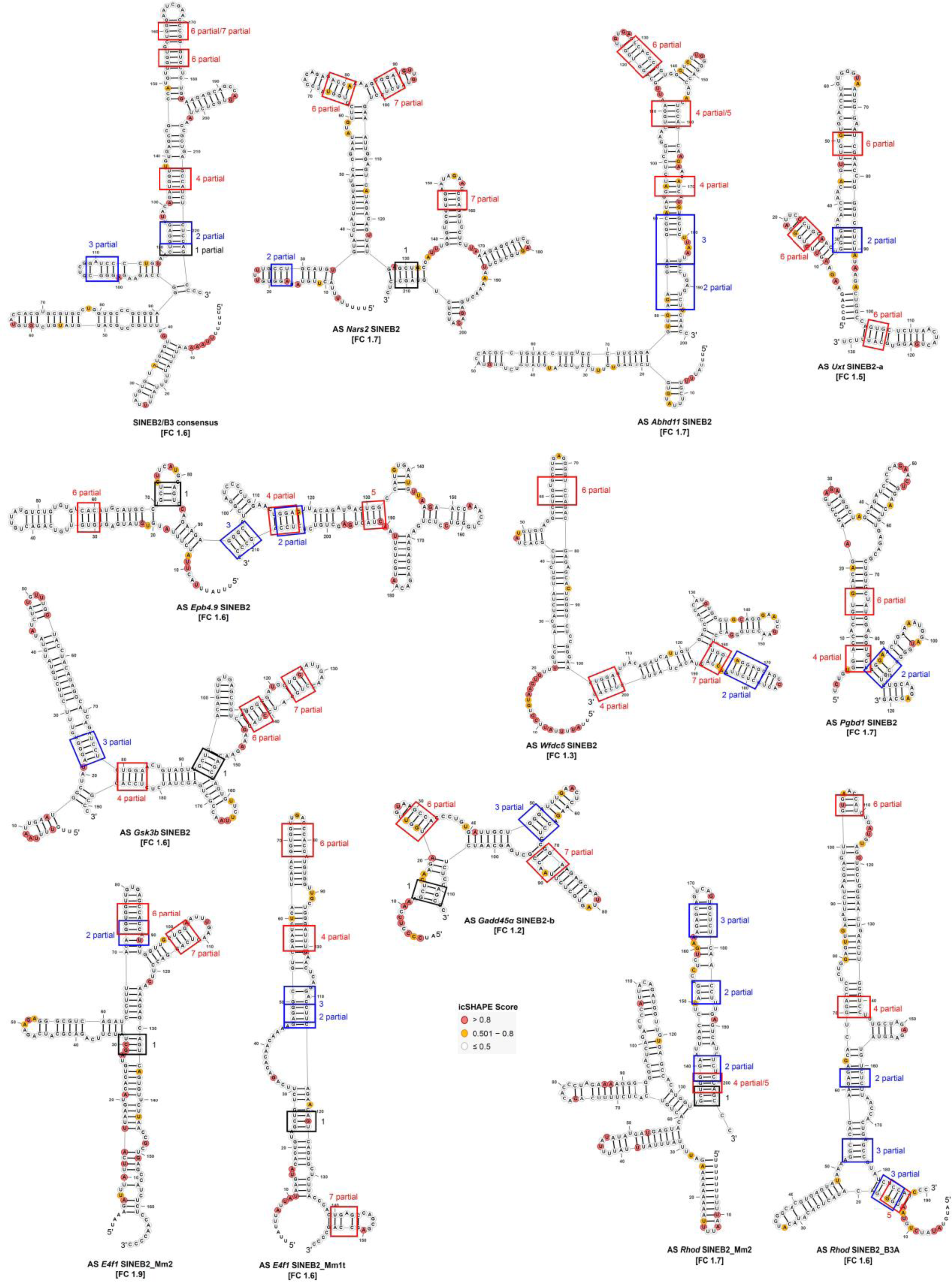
icSHAPE 2D structure models of mouse SINEB2 RNAs from natural antisense transcripts that are tested as effector domain in SINEUPs. Regions that match the SINEUP motifs are marked by squares with their corresponding motif number. Blue square, AGG type motif; red square, UGG type motif. FC, GFP protein fold-change induced by the respective SINEB2. Nucleotide color indicates normalized icSHAPE reactivity score: red, ≥ 0.8; yellow, from above 0.5 to 0.8; white encircled with grey, ≤ 0.5; bases not encircled, no icSHAPE data available. AS, antisense.

**Extended Data Figure 5.**
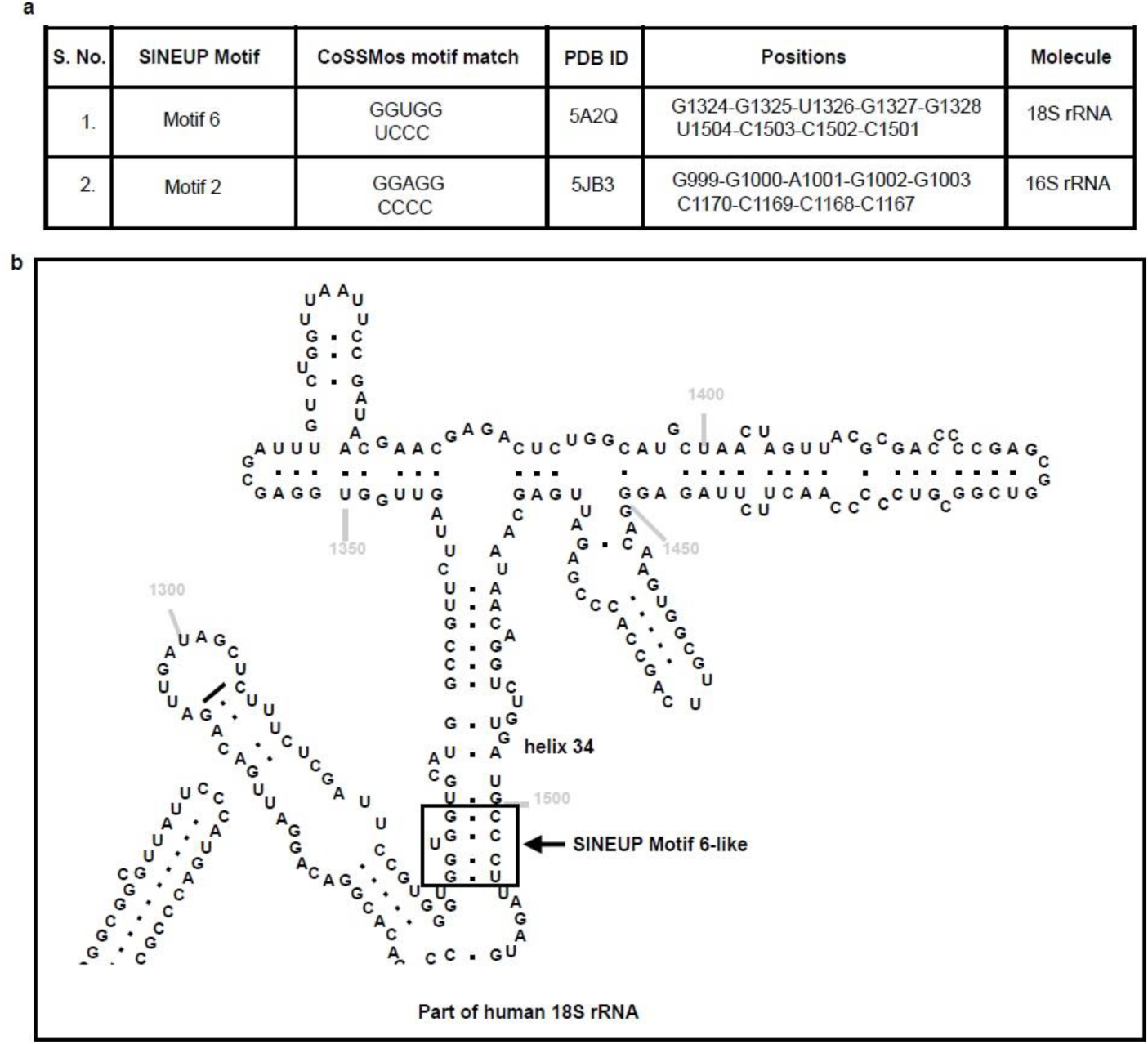
Similarity of SINEUP motifs to rRNA 3D structure motifs. (**a**) Hits found in the Characterization of Secondary Structure Motifs database for 3D motifs that match the SINEUP structure motifs. (**b**) The region similar to SINEUP motif 6 in helix 34 of human 18S rRNA is framed in black and marked with an arrow.

**Extended Data Figure 6.**
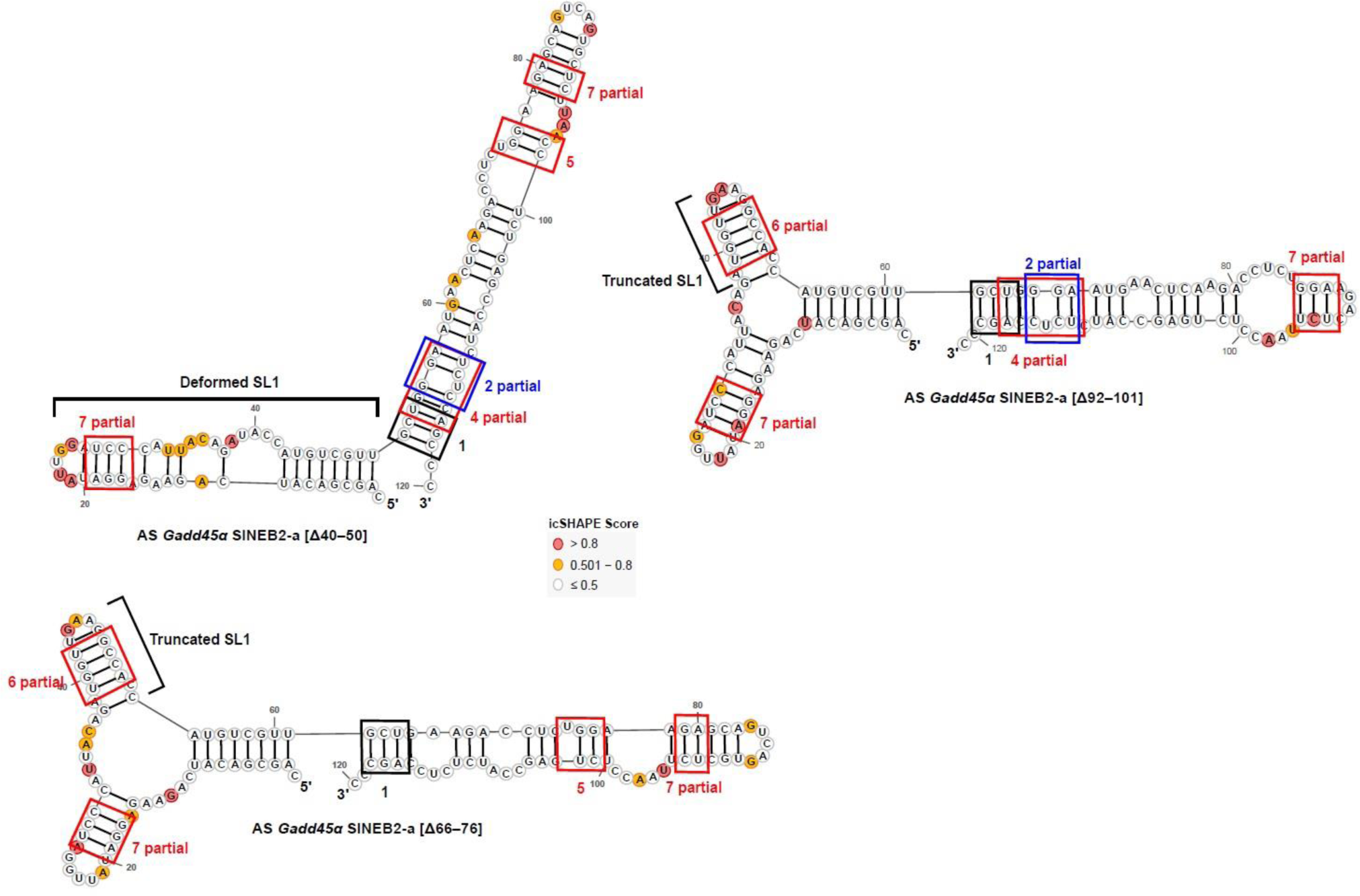
SINEUP structure motifs in functionally inactive antisense (AS) *Gadd45α* SINEB2-a deletion mutants. icSHAPE derived structures of AS *Gadd45α* SINEB2-a deletion mutants [Δ40– 50], [Δ66–76], and [Δ92–101]. Regions that match conserved SINEUP structure motifs are marked by different colors and numbers. Δ = deletion. Nucleotide color indicates normalized icSHAPE reactivity score: red, ≥ 0.8; yellow, from above 0.5 to 0.8; white encircled with grey, ≤ 0.5; bases not encircled, no icSHAPE data available. AS, antisense.

**Extended Data Figure 7.**
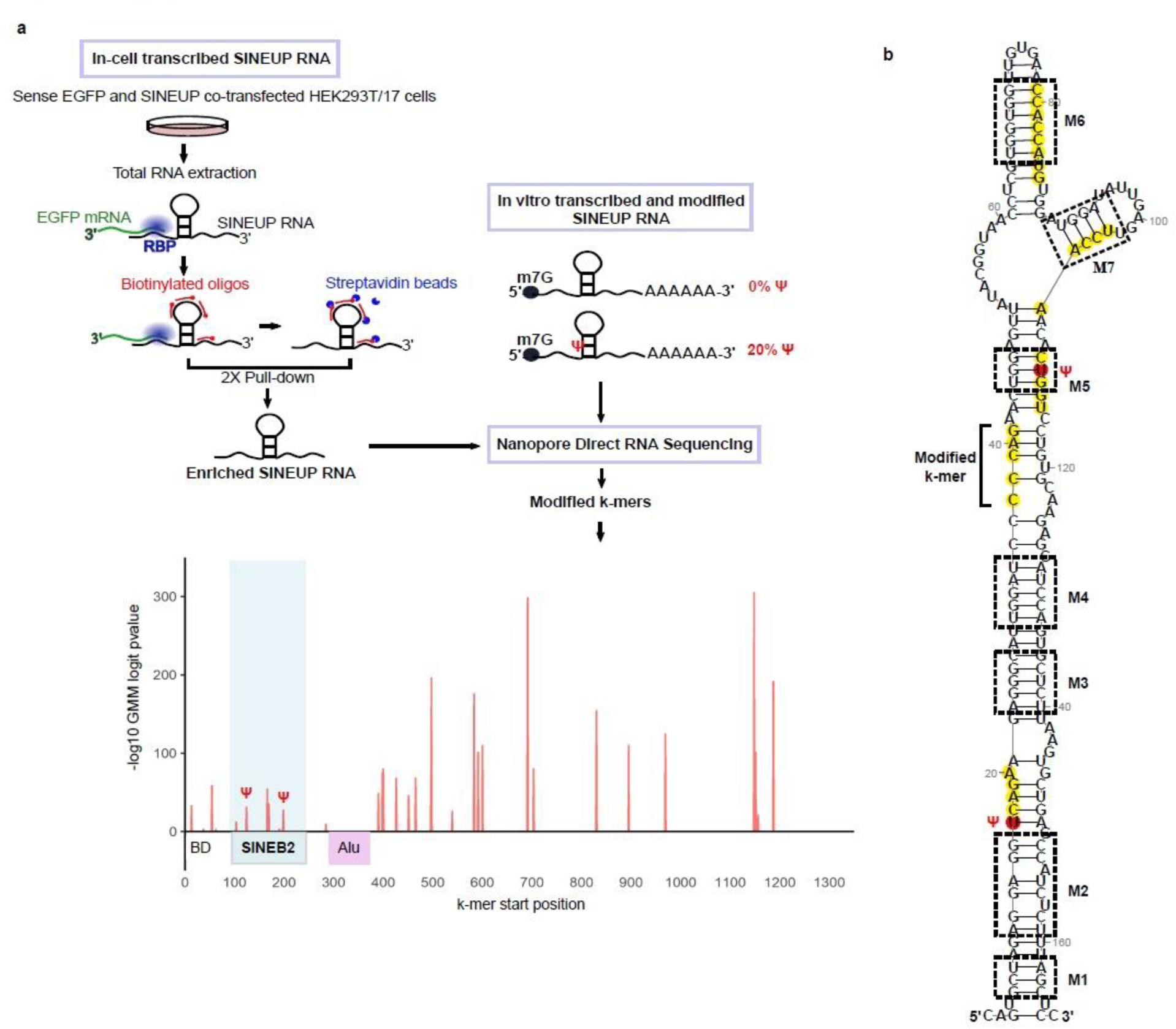
Discovery of modified bases in SINEB2 RNA. (**a**) An overview of SINEUP RNA modification analysis. SINEUP-GFP RNA (containing antisense Uchl1 SINEB2) transcribed in vitro was modified with 20% pseudouridine (Ψ), and a non-modified (0% Ψ) sample was used as control. In parallel, SINEUP-GFP and sense EGFP plasmids were co-transfected in HEK293T/17 cells, then SINEUP RNA that was transcribed in-cell was captured and enriched using specific biotinylated oligo probes and pull-down (twice) on streptavidin-coated magnetic beads. Purified samples of SINEUP-GFP RNA transcribed in-cell or transcribed and modified in vitro were sequenced by Nanopore direct RNA sequencing. Profiles of non-modified (0% Ψ) and modified SINEUP-GFP RNA transcribed in vitro were compared with that transcribed in-cell, and k-mers corresponding to the modified positions were identified in the in-cell transcripts. The red peaks in the graph mark the position of modified k-mers on the SINEUP transcript. The SINEB2 region is shaded in blue, and verified Ψ sites are marked. BD, binding domain. (**b**) Modified k-mers overlaid on 2D structure of antisense *Uchl1* SINEB2. k-mers are highlighted in yellow, Ψ sites are in red. SINEUP motifs are marked in black dashed squares with motifs 1–7 labelled as M1–M7.

**Extended Data Figure 8.**
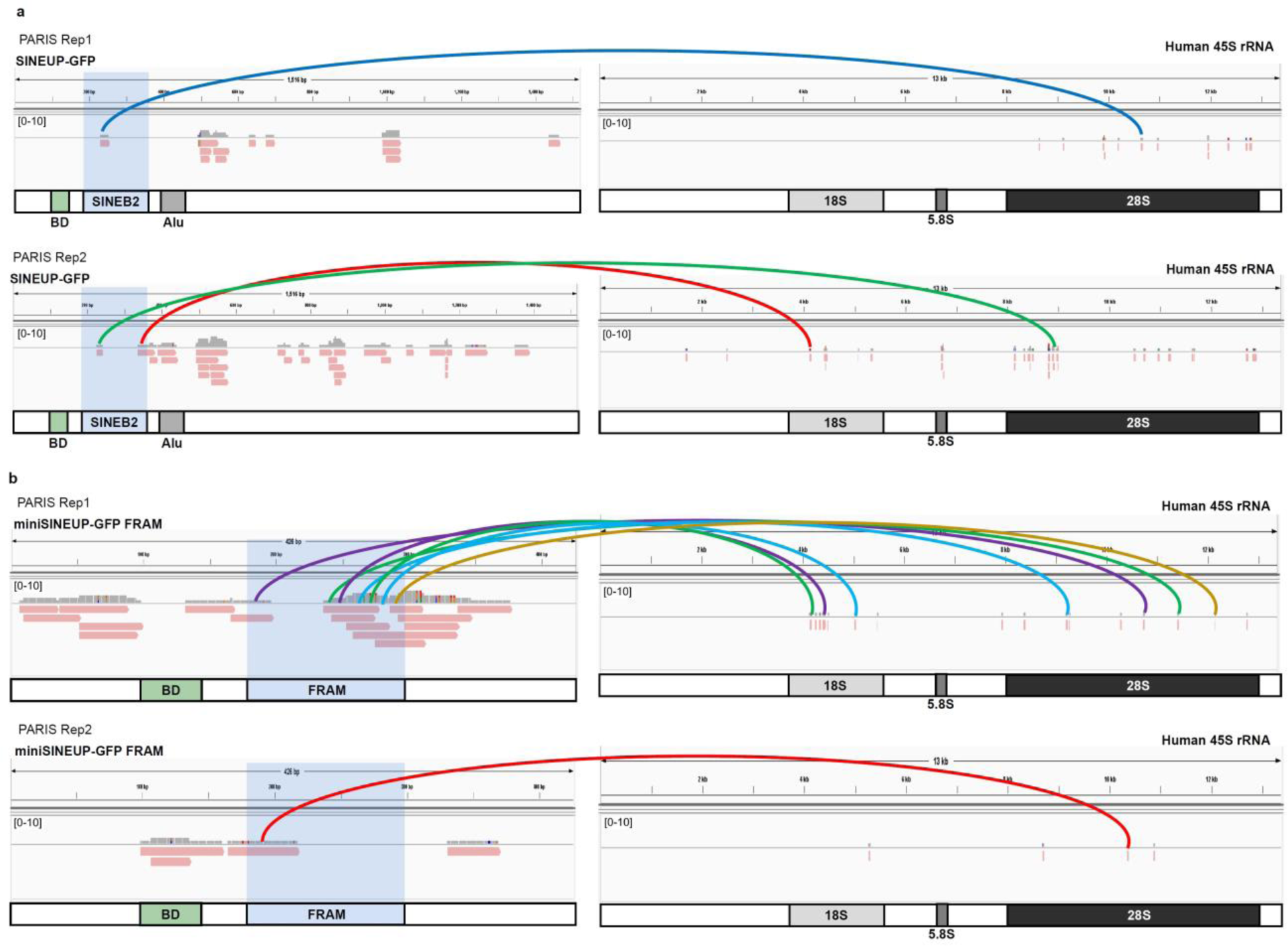
SINEUP-rRNA duplex reads in PARIS sequencing data. (**a**) SINEUP-GFP (of antisense *Uchl1* SINEB2) RNA duplex reads with 18S, 5.8S, and 28S rRNA. Two biological replicates of the PARIS experiment are shown as separate tracks. SINEB2-rRNA interactions shown in Fig. 4 are indicated by respective colored arcs. (**b**) miniSINEUP-GFP FRAM RNA-rRNA duplex reads. PARIS rep1 and rep 2 are two biological replicates. The FRAM RNA-rRNA interactions displayed in Fig. 5 are marked by arcs in their respective colors.

**Extended Data Figure 9.**
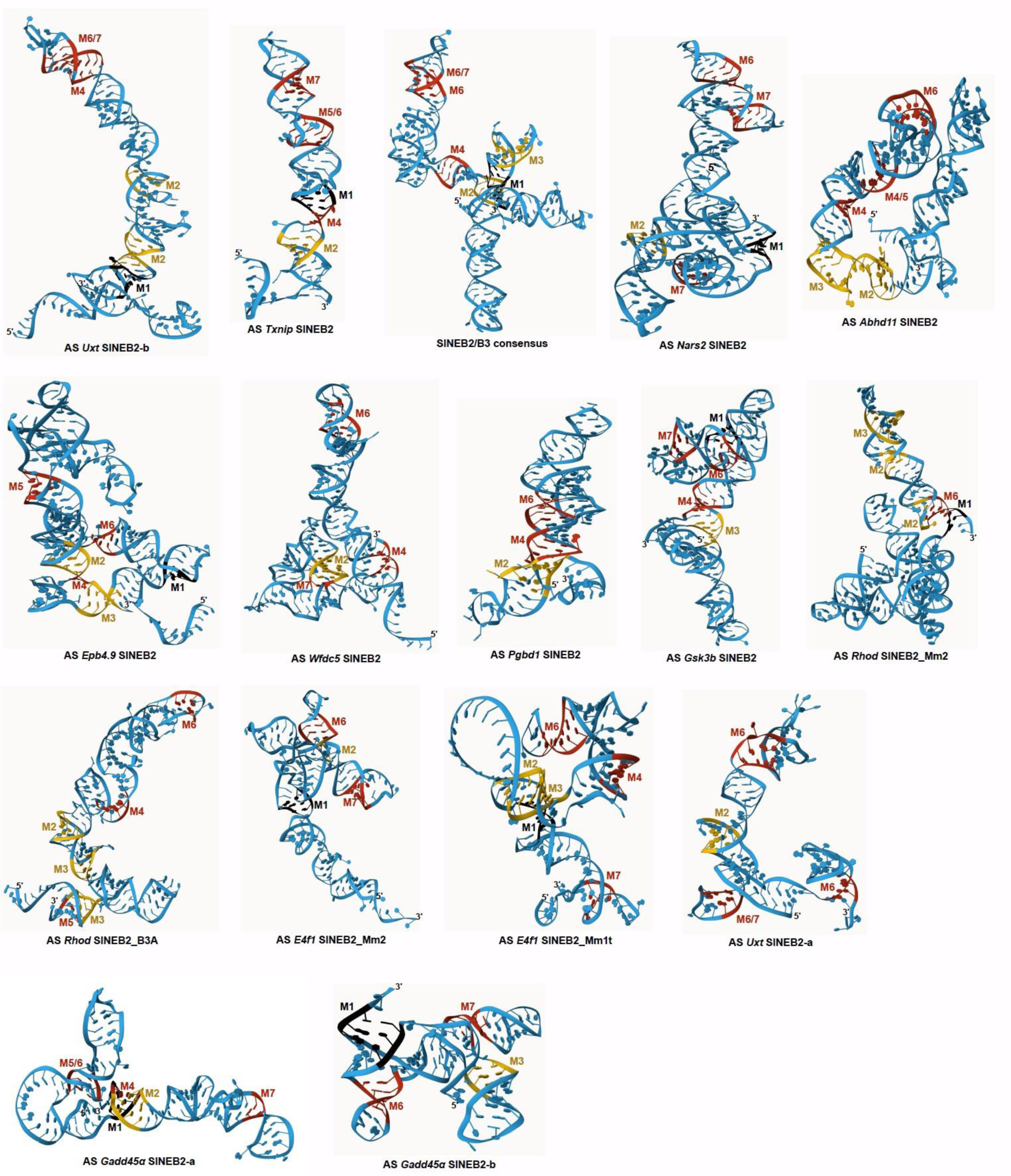
SINE RNA predicted 3D structure models based on in-cell 2D structures of mouse SINEB2 RNAs. SINEUP structure motifs are denoted from M1 to M7. AS: antisense.

**Extended Data Figure 10.**
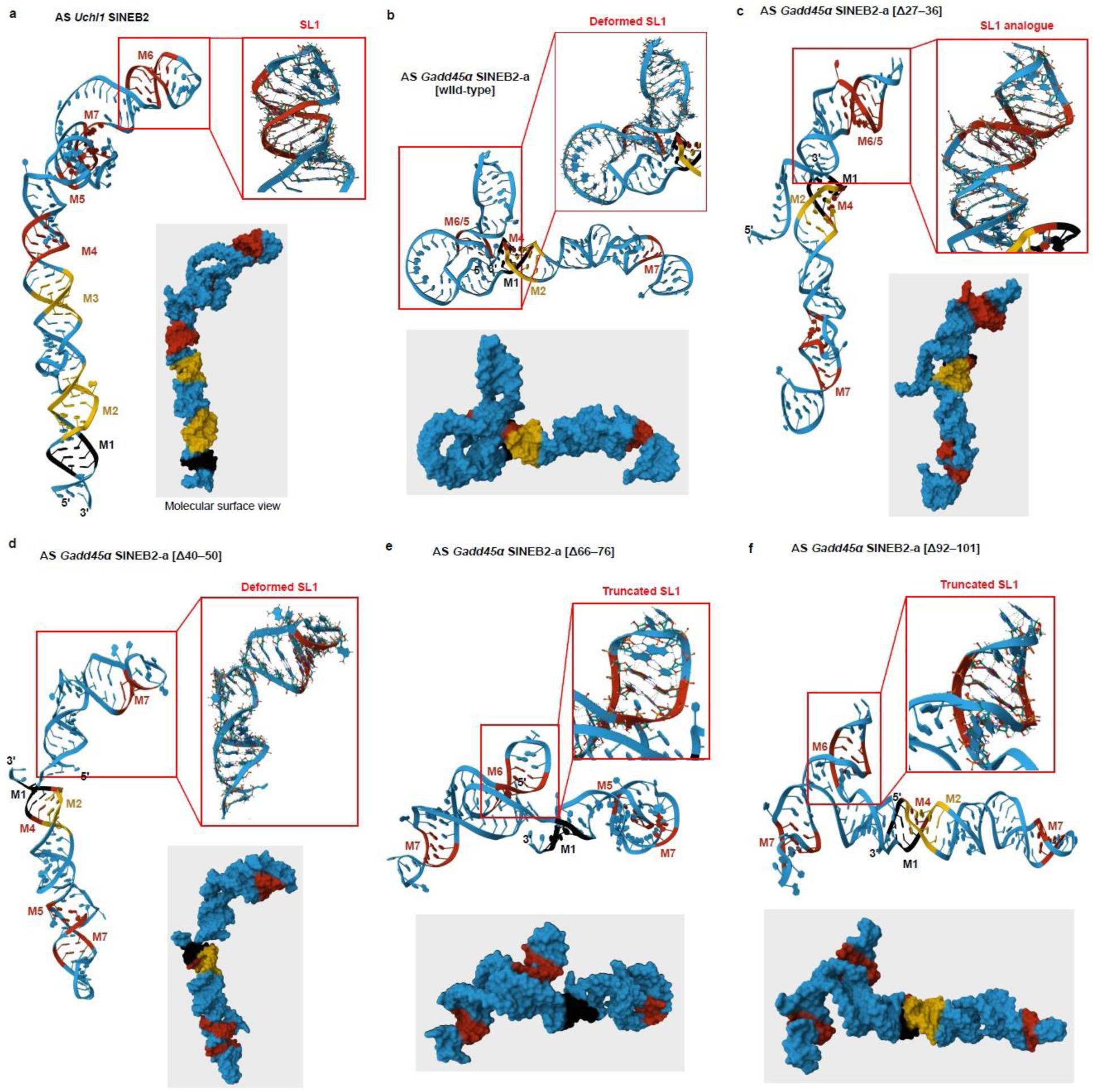
Comparison of icSHAPE data-driven predicted 3D structure models of wild-type AS *Gadd45α* SINEB2-a and its deletion mutants with functional AS *Uchl1* SINEB2. (**a**) AS *Uchl1* SINEB2 3D model predicted from whole cell icSHAPE data. Predicted 3D models with enlarged view of the corresponding SL1 regions for AS *Gadd45α* SINEB2-a (**b**) wild-type (functionally inactive), its (c) functionally active deletion mutant [Δ27–36], and non-functional mutants (d) [Δ40–50], (e) [Δ66–76], and (f) [Δ92–101]. SINEUP structure motifs M1–M7 are marked. Red frame shows zoomed-in view of the region corresponding to SL1. The molecular surface views of the 3D models are shaded in gray. Δ = deletion.

## Notes

### Summary of Updates

New icSHAPE-derived structures for SINEB2 deletion mutants have been added in Figure 2 and new Extended Data Figure 6. 3D structure models for SINEB2 deletion mutants have been added in new Extended Data Figure 10. The results section has been revised extensively.

